# Cytoplasmic mRNA decay by the anti-viral nuclease RNase L promotes transcriptional repression

**DOI:** 10.1101/2025.07.08.663778

**Authors:** Xiaowen Mao, Sherzod Tokamov, Felix Pahmeier, Azra Lari, Jinyi Xu, Eva Harris, Britt Glaunsinger

## Abstract

Ribonuclease (RNase) L is an antiviral factor that promiscuously degrades viral and cellular RNA in the cytoplasm. This results in extensive translational reprogramming, altering mRNA processing and export. Here, we reveal that another major consequence of cytoplasmic RNase L activity is the repression of nascent RNA synthesis in the nucleus. This is not associated with altered nuclear RNA stability but instead results from transcriptional repression. For RNA polymerase II, repression is primarily associated with reduced occupancy of serine 2-phosphorylated polymerase in gene bodies, indicating an elongation defect. Prominent among the transcriptionally downregulated loci are immune-related genes, supporting a role for RNase L in tempering innate immune and inflammatory responses. RNase L activation also caused disruption of nucleoli and reduced RNA polymerase I and III transcription. Crosstalk between RNA decay and transcription thereby contributes to the large-scale modulation of gene expression in RNase L-activated cells.

## Introduction

Ribonuclease (RNase) L is an important player in the innate immune response triggered by double-stranded RNA (dsRNA), a pathogen-associated molecular pattern (PAMP) commonly associated with viral infections. RNase L dimerizes and is activated by 2’-5’-oligo(A), a second messenger produced by the oligoadenylate synthetases (OAS) 1-3 upon dsRNA binding. Once activated, RNase L promiscuously cleaves cytoplasmic RNA, including viral RNAs and host rRNA, mRNA, and tRNA, at sites defined by UN^N ^1^. RNase L was initially hypothesized to suppress translation through its ability to cleave rRNA and tRNA. However, rRNA and tRNA cleavage occurs after translation remodeling, and ribosomes with RNase L-cleaved rRNAs can still support translation ^2–4^. Thus, widespread cleavage of mRNA most likely causes the prominent decrease in cellular translation ^2,4,5^.

In addition to suppressing viral gene expression, RNase L-induced gene expression reprogramming both positively and negatively modulates innate immune responses ^2,4–12^. RNAse L promotes virus-induced IFNβ production in mice and in cells where RNase L basal expression is low, but suppresses IFNβ production in cells with higher basal RNase L and OAS expression, perhaps to promote tissue protection ^10,13^. Additionally, loss-of-function mutations in the OAS-RNase L pathway have been linked to autoimmune diseases, including some cases of multisystem inflammatory syndrome in children following SARS-CoV-2 infection ^9^. These observations suggest that RNase L activity may dampen immune gene expression to prevent overactivation of the innate immune system. However, some mRNAs encoding antiviral factors are at least partially resistant to RNase L and may be preferentially translated due to the liberated translational resources, ensuring the execution of an antiviral response ^2,4,5^. Much remains to be learned about the contexts governing the complex roles of RNase L in immune regulation.

While RNA cleavage is a prominent phenotype associated with RNase L, several recent findings reveal additional mechanisms contributing to its impact on gene expression. These include inhibiting mRNA export ^6^, altering mRNA processing ^7^, activating ribotoxic stress responses ^8,11,14^, and activating RIG-I/MDA5-MAVS pathways through the production of small RNA cleavage fragments ^10,13^. Each of these phenotypes relies on RNase L catalytic activity, suggesting that they occur in response to the widespread RNA cleavage.

Several oncogenic gamma-herpesviruses also engage broad-acting mRNA-targeting nucleases, whose activity, like that of RNase L, can lead to alterations in mRNA processing and export ^15–18^. Notably, viral cleavage of cytoplasmic mRNA also induces large-scale dampening of RNA polymerase II (Pol II) transcription ^19–21^. Similarly, widespread cytoplasmic mRNA decay by the cellular exonuclease polyribonucleotide nucleotidyltransferase 1 (PNPT1), which is activated during early apoptosis, leads to subsequent Pol II transcriptional repression ^22–24^. These findings indicate that there is a positive feedback loop between mRNA synthesis and degradation under conditions of accelerated mRNA decay, which we hypothesized might also occur in response to RNase L activation.

Here, we reveal that activated RNase L represses RNA synthesis. This includes interferon-stimulated genes and other immune genes, suggesting that its impact on RNA biogenesis may help temper inflammatory responses. The reduction of newly synthesized RNA is not due to accelerated RNA decay in the nucleus but instead is triggered indirectly by RNase L-induced RNA degradation in the cytoplasm. This causes a reduction in transcription by Pol I, II, and III, accompanied by alterations in nucleolar and nuclear speckle morphology. The primary alteration to Pol II occupancy is a reduction in the serine 2-phosphorylated version in the gene body and near transcription termination sites, suggestive of an elongation defect. The repressive signal involves the transport of proteins into the nucleus, as blocking importin β-mediated nuclear import rescues RNA synthesis in cells with activated RNase L. Thus, crosstalk between the cytoplasm and nucleus promotes broad transcriptional repression upon RNase L activation, which contributes to the reshaping of the gene expression landscape in cells following innate immune stimulation.

## Results

### RNase L activation broadly reduces the abundance of newly synthesized RNA in the nucleus

Based on observations of positive feedback between RNA decay and synthesis ^19,24^, we sought to determine how RNase L impacts the abundance of newly synthesized RNA in the nucleus. We first designed an assay to quantify the total pool of newly transcribed RNA in response to RNase L activation. We activated RNase L by transfecting human A549 cells with the double-stranded RNA mimic poly(I:C) for 4 hours, then added the uridine analog ethynyl uridine (EU) for 30 minutes to label nascent transcripts. The EU-containing RNA was coupled to a Cy5 fluorophore by click chemistry and then visualized by flow cytometry or imaging (Figure 1A). We confirmed that the labeled RNA remained largely nuclear during the 30-minute pulse (Figure S1A-B), which is important because, upon export to the cytoplasm, RNA is rapidly degraded by RNase L.

**Figure 1.**
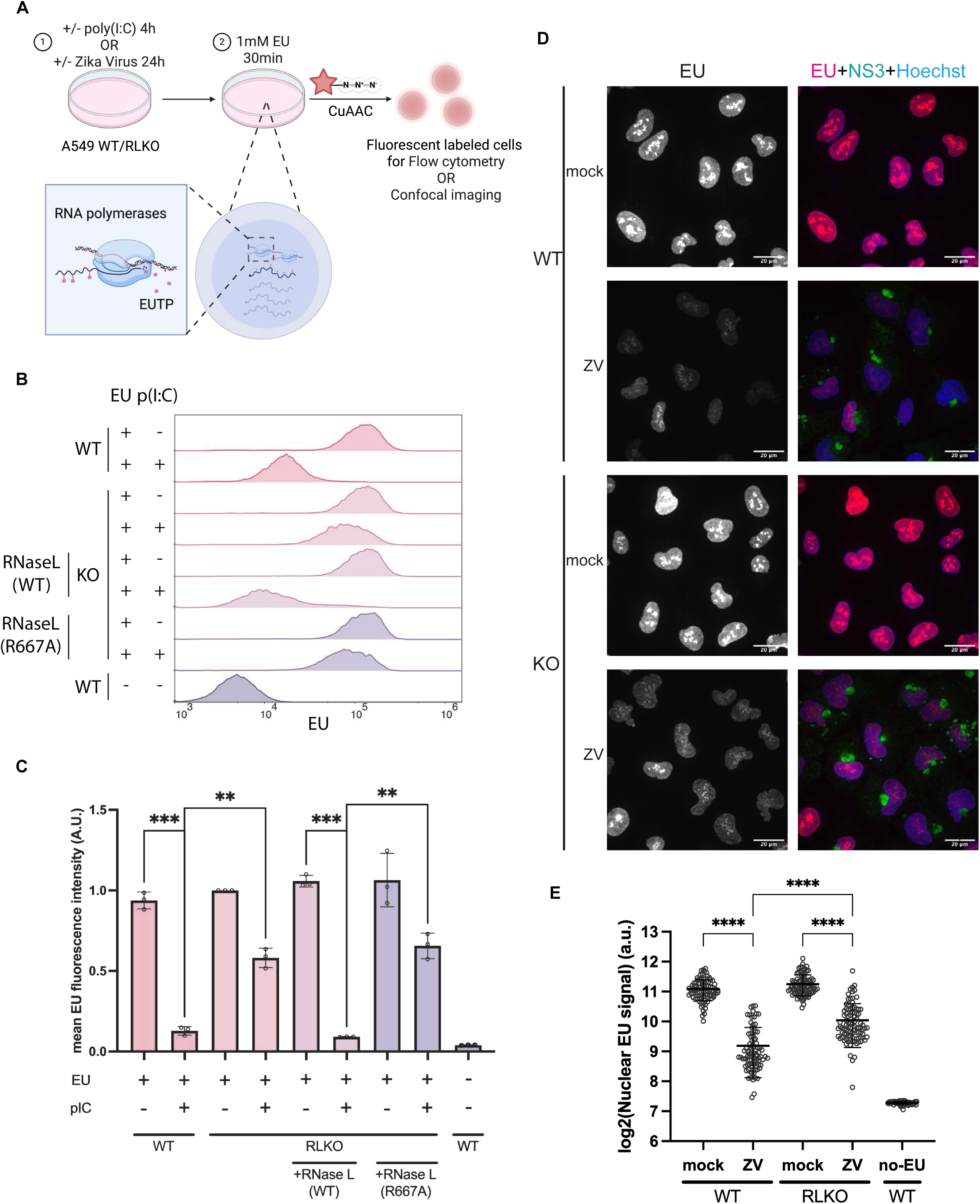
RNase L-mediated mRNA degradation represses nascent RNA accumulation. (A) Diagram showing the experimental setup for nascent RNA measurement. (B) Flow cytometry histograms of A549 WT or RNase L knockout (RLKO) cells with the indicated treatment from one representative biological replicate (of 3 independent biological replicates). (C) Quantification of flow cytometry data from 3 independent biological replicates. Each dot represents the median fluorescence intensity of the sample with the indicated treatment from one biological replicate, normalized to the median fluorescence intensity of mock-transfected RLKO cells. Bars represent mean ± SD. P-values were calculated using a two-sided Welch’s *t*-test. (D) Representative immunofluorescence images (max intensity projection, 3 independent biological replicates) of A549 WT or RLKO cells infected with Zika virus (MOI=5) at 24 hpi. Cells were stained with mouse anti-NS3 antibody to indicate infected cells, and Hoechst 33342 to mark nuclei. (E) Quantification of nuclear EU signal in 1 representative replicate of 3 independent biological replicates in *(D)*. Each dot represents the mean nuclear EU fluorescence intensity of one individual nucleus. Bars represent mean ± SD. P-value was calculated using Welch’s *t* test.

We observed a striking reduction in EU incorporation in poly(I:C)-treated WT A549 cells compared to control mock-treated cells (Figure 1B, C). In contrast, poly(I:C) treatment of RNase L knock-out (RLKO) A549 cells resulted in only a modest reduction in EU signal (Figure 1B, C), suggesting that RNase L is the primary driver of the loss of newly transcribed RNA in the nucleus. Furthermore, nascent RNA depletion was observed in A549 RLKO cells complemented with WT RNase L but not those complemented with a catalytically dead RNase L mutant R667A (Figure 1B, C), indicating that this effect required RNase L catalytic activity. This phenotype was detected as early as 2 hours after poly(I:C) treatment and grew progressively more pronounced over time (Figure S1C). It was also apparent in WT but not RLKO cells treated with a synthetic version of the RNase L-activating second messenger molecule 2-5A (pA(_2-5_A)_4_) (Figure S1D). Poly(I:C) transfection also reduced the EU signal in immortalized murine bone marrow-derived macrophages (iBMDM), and this effect was mitigated upon siRNA-mediated depletion of RNase L (Figure S1E-G). Although RNase L activation can induce caspase-dependent apoptosis ^12,25^, the irreversible pan-caspase inhibitor z-VAD(Ome)-FMK did not prevent the loss of EU signal (Figure S1H), indicating that repression of nascent RNA does not require caspase activation.

To determine whether these observations occur during viral infection, we performed a 30-minute EU pulse during Zika virus infection, which is a potent activator of RNase L ^26–28^. Indeed, Zika virus infection at an MOI of 5 for 24 hours resulted in a significant reduction in EU-labeled RNA in WT A549 cells compared to RLKO cells, as visualized by immunofluorescence (Figure 1D-E, S1I). A portion of the Zika virus-induced reduction in nascent RNA was RNase L-independent (Figure 1E), which may stem from other effects induced by dsRNA and/or Zika virus capsid protein ^29^. Collectively, these data indicate that RNase L-mediated mRNA degradation represses nascent RNA synthesis in multiple cell types and in response to multiple activating stimuli.

### RNase L reduces both nascent rRNA and pre-mRNA accumulation

EU is incorporated into transcripts by all three classes of RNA polymerases, with RNA polymerase I (Pol I) and Pol II transcripts each accounting for approximately half of the total signal during a short pulse ^30^. Thus, the loss in EU signal caused by RNase L activation is likely due to the reduction of both nascent Pol I and Pol II transcripts. To comprehensively investigate the nascent transcripts affected by RNase L activation, we sequenced the newly transcribed pool of RNA following a 10-minute pulse labeling with another uridine analog, 4-thiouridine (4sU). After adding labeled and unlabeled spike-in controls for normalization, the 4sU-incorporated RNA was purified away from the pre-existing pool of RNA by biotinylation of the 4sU and purification via streptavidin beads (Figure 2A). We confirmed >30-fold enrichment of the 4sU-labeled over the unlabeled spike-in RNA in our sequencing data and confirmed 4sU RNA enrichment by RT-qPCR (Figure S2A-B).

**Figure 2.**
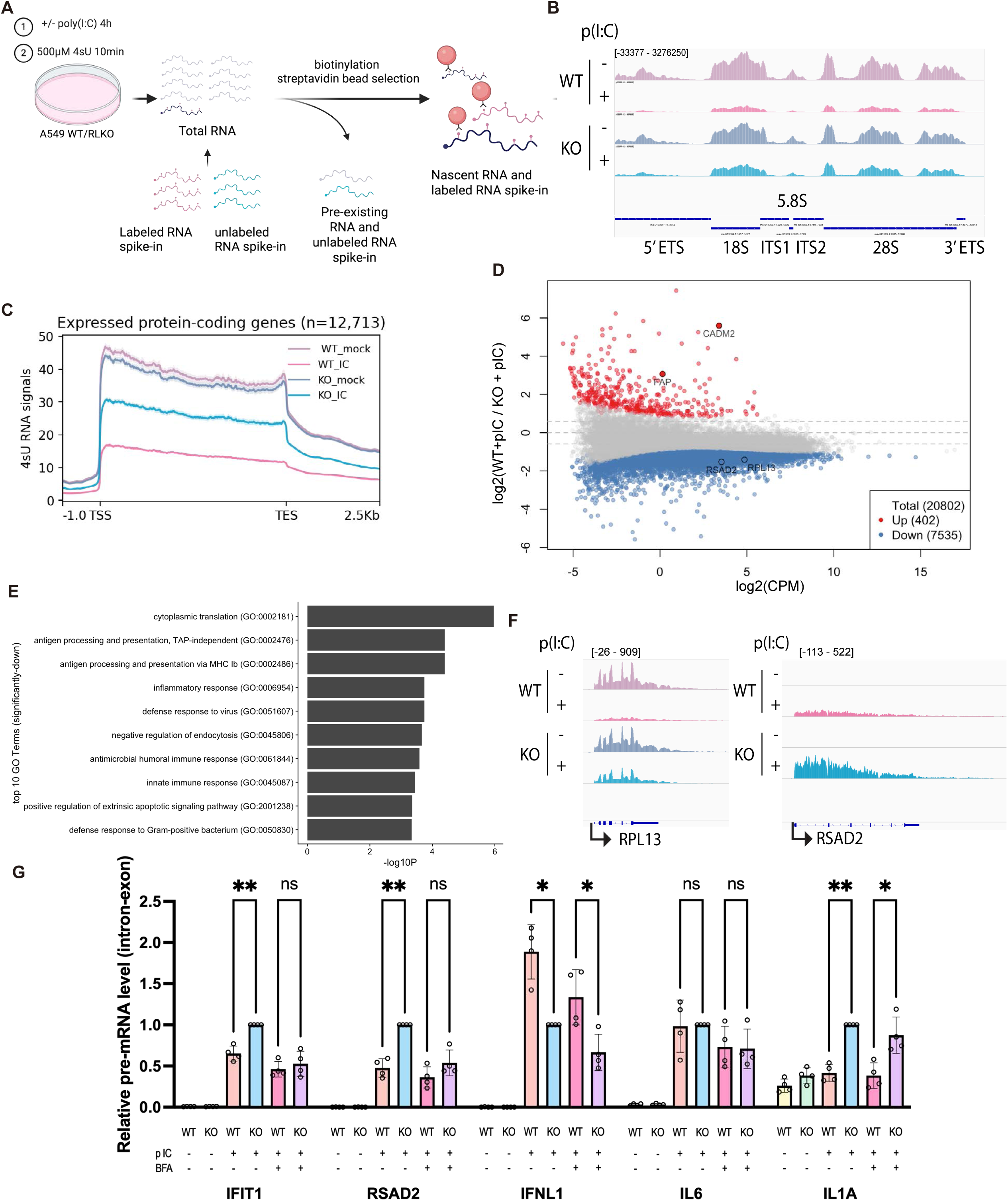
RNase L reduces nascent rRNA and pre-mRNA. (A) Diagram summarizing the 4sU-seq experiment. (B) IGV tracks showing 4sU-seq coverage at rDNA repeat. Tracks represent the average coverage of 3 independent biological replicates. Coverage depth is shown in brackets in the upper track, with a positive value indicating coverage in the positive strand direction, and a negative value indicating coverage in the negative strand direction. (C) Metagene analysis of 4sU-seq signal coverage over expressed protein-coding genes (-1kb from TSS to +2.5kb from TES, n=12,713). Lines represent the average coverage of 3 independent biological replicates. (D) Differential expression analysis of 4sU-seq comparing WT cells transfected with poly(I:C) to RLKO cells transfected with poly(I:C). Each gene was plotted with the log2(fold change) value on the y-axis and the log2 of the mean expression level, expressed in counts per million (CPM), on the x-axis. Significantly-up (red) and significantly-down (blue) are defined as adjusted *p*-value < 0.05 and log2(fold change) > 0.585 or <-0.585, respectively. Dashed lines denote log2(fold change) of -0.585, 0, and 0.585. (E) GO-term enrichment analysis of down-regulated genes identified in *(D)*. (F) IGV tracks showing 4sU-seq signal coverage on RPL13, an example of ribosomal protein genes, and RSAD2, an example of immune genes. Coverage depth is shown in brackets in the upper track, with a positive value indicating coverage in the positive strand direction, and a negative value indicating coverage in the negative strand direction. (G) Pre-mRNA level of immune genes in WT or RLKO cells transfected with poly(I:C), with or without treatment of brefeldin A (BFA). Pre-mRNA level is determined by RT-qPCR using intron-exon spanning primer pairs, normalized to 18S. *, p< 0.05; **, p<0.01; ***, p< 0.001; ns, not significant.

Initial rRNA filtering revealed that the rRNA reads accounted for no more than 56% of the total reads, which aligns with expectations ^31^. After alignment to the rDNA repeat and spike-in normalization, we observed a decrease in 4sU-seq signal coverage on the entire rDNA repeat in WT cells transfected with poly(I:C), which was less prominent in RLKO cells (Figure 2B). We then aligned the remaining reads to the hg38 reference genome and performed a metagene analysis on protein-coding genes. In agreement with the EU flow cytometry data, we observed a global decrease in nascent mRNA transcripts in poly(I:C) treated WT cells relative to mock treated WT cells, as well as in WT cells treated with poly(I:C) compared to RLKO cells treated with poly(I:C) (Figure 2C). We also observed a prominent decrease in several Pol III-transcribed genes (Figure S2C). Differential gene expression analysis of protein-coding and non-coding genes transcribed by all three RNA polymerases revealed a widespread decrease on a gene-by-gene basis, with 36.2% of genes significantly downregulated (Figure 2D). Thus, RNase L plays a major role in repressing the accumulation of nascent rRNA, noncoding RNA, and mRNA.

Fewer genes were significantly upregulated in an RNase L-dependent manner, and manual inspection of these genes revealed that their apparent upregulation is largely explained by one of two situations. The first category contains genes that are differentially expressed in these cells even in the absence of poly(I:C) treatment, indicating they are the result of minor genetic/epigenetic differences between the parental WT cell line and the clonal RLKO cell line (Figure S2D-E). The second category includes genes whose apparent upregulation is instead the result of read-through transcription from upstream highly expressed genes (Figure S2F). Thus, RNase L activation predominantly causes repression rather than activation of nascent RNA.

Notably, gene ontology analysis of the downregulated genes revealed an enrichment of genes involved in cytoplasmic translation and the immune response (Figure 2E-F). The translation term is likely due to the reduction in ribosomal protein-encoding mRNAs, which are known targets of RNase L ^32^, suggesting a coordinated response with rRNA transcription inhibition to reduce ribosome biogenesis ^33^. The abundance of immune-related categories suggests that RNase L-induced suppression of nascent RNA contributes to its impact on the immune response. We validated this result by performing RT-qPCR on selected immune transcripts in total RNA samples using intron-exon spanning primer pairs to measure pre-mRNA. Indeed, interferon-stimulated genes (ISGs), including IFIT1, RSAD2, OAS1, and OASL, displayed decreased pre-mRNA levels in WT cells compared to RLKO cells stimulated with poly(I:C) (Figure 2G, S2G). In contrast, IFNL1 has previously been shown to undergo intron retention upon RNase L activation, and we observed this expected increase (and a similar trend for ISG15) in WT relative to RLKO cells (Figure 2G).

We next considered the extent to which the decrease in ISG pre-mRNA was due to a reduction in interferon (IFN) signal amplification versus a cell-autonomous effect, as we observed decreased levels of both nascent IFNB1 and IFNL1. This is also relevant because degradation and/or export inhibition of interferons and other cytokine cytoplasmic mRNA by RNase L should reduce the level of their encoded secreted proteins, thereby dampening their ability to induce ISG expression in neighboring cells via paracrine signaling. To evaluate this for ISGs, we treated cells concurrently with poly(I:C) and brefeldin A (BFA), which collapses Golgi-ER retrograde transport and inhibits protein secretion ^34^. BFA reduced the difference in nascent mRNA levels between WT and RLKO cells for most of the ISGs we tested, including IFIT1, RSAD2, OAS1, and OASL, confirming the impact of RNase L activity on signal amplification of the interferon pathway (Figure 2G, S2G). However, the difference in IL1A nascent mRNA levels between WT and RLKO cells remained upon BFA treatment (Figure 2G), suggesting that the repression of nascent RNA can independently influence some immune genes in a cell-autonomous manner.

### Reduction of nascent RNA is not due to accelerated nuclear RNA degradation

RNase L is primarily described as a cytoplasmic nuclease ^7,35^; however, even a small amount of active RNase L in the nucleus could lead to rapid depletion of pre-mRNA. To directly test whether RNase L activation increases degradation of nascent RNA in the nucleus, we measured nascent RNA turnover using an EU pulse chase experiment. We pulsed poly(I:C)-transfected cells with EU 1 hour after transfection, a time point when EU incorporation is minimally affected by RNase L activation (Figure S1C, S3A). We then chased with excess uridine over a 3-hour time course and measured the decay of the labeled RNA. Notably, the rate of RNA decay was similar in A549 WT and A549 RLKO cells, indicating that RNase L does not accelerate nascent RNA decay in the nucleus (Figure 3A). We confirmed that the transfection conditions were sufficient to induce RNase L-specific reduction of nascent RNA (Figure S3A-B, see the “pulse last” sample) and that our measurements largely captured nuclear RNA (Figure S3C-D).

**Figure 3.**
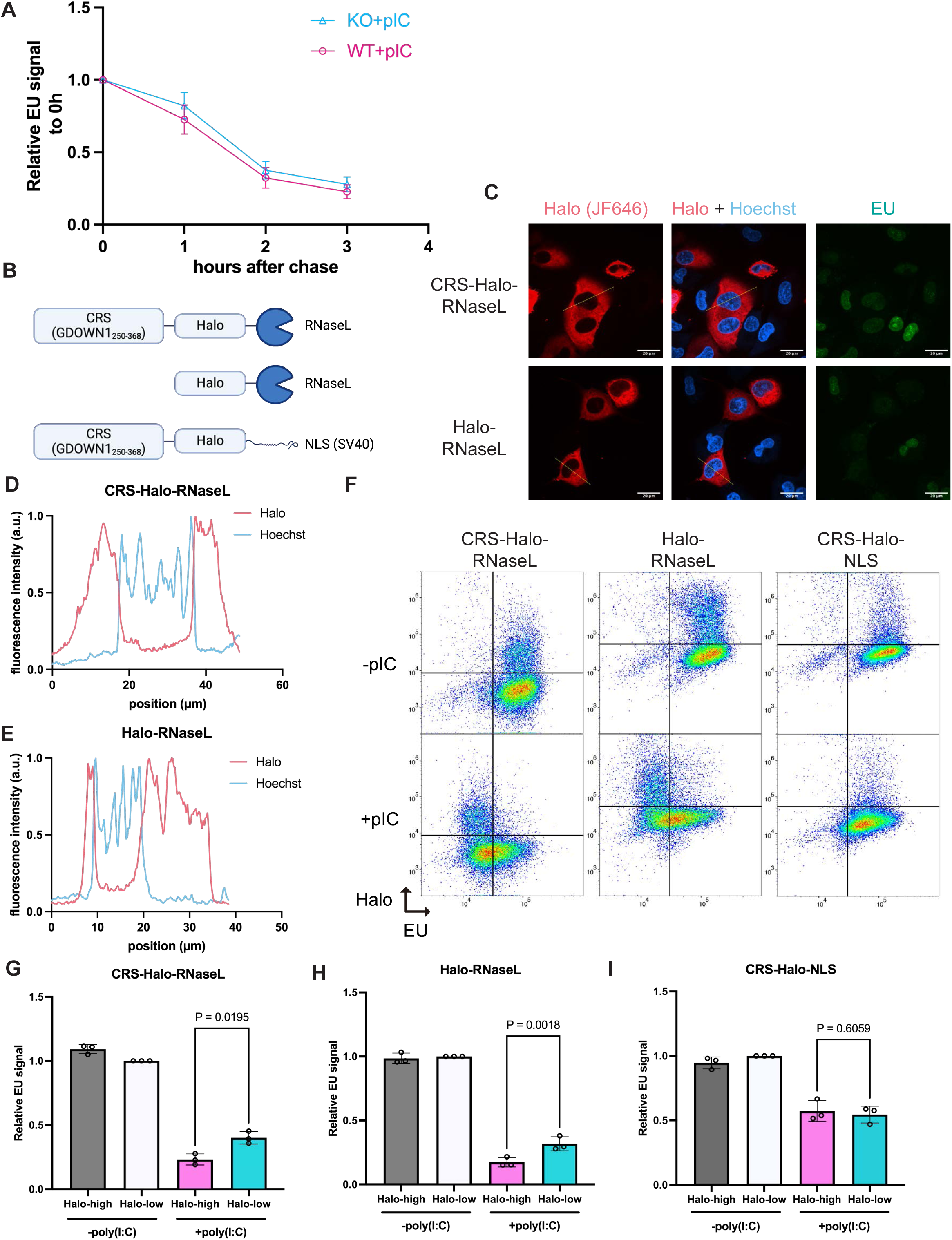
Reduction of nascent RNA is not due to accelerated nuclear RNA degradation by RNase L. (A) Quantification of EU signal intensity after uridine chase for durations indicated. Each point represents the average of median EU fluorescence intensity from 3 independent biological replicates, normalized to 0h chase. (B) Diagrams of the constructs used. (C) Representative images of the localization of tagged RNase L. The tagged RNase L was visualized by halo tag labeling with JF646 halo dye. A single z plane is shown per construct. (D) Line profile of CRS-Halo-RNase L shown in *(C)*. (E) Line profile of Halo-RNase L shown in *(C)*. (F) Flow cytometry scatter plot of A549 RLKO cells nucleofected with the indicated constructs, with or without poly(I:C) transfection. Data is from 1 representative replicate of 3 independent biological replicates. (G-I) Quantification of EU signals in *(F)*. The Halo intensity is gated at locations shown in (F) and separated into 2 populations. Each dot represents the median fluorescence intensity, normalized to the median fluorescence intensity of mock-transfected RLKO cells nucleofected with the constructs indicated. Bars represent mean ± SD. P-values were calculated using a ratio paired *t*-test.

As an orthogonal way of confirming that the cytoplasmic fraction of RNase L is sufficient to reduce nascent RNA in the nucleus, we complemented the RNase L knockout cells with a Halo-tagged version of RNase L fused to a cytoplasmic retention signal (CRS) (Figure 3B) derived from GDOWN1 (amino acid 250-368) ^36^. This CRS promotes cytoplasmic retention by both CRM1-dependent and -independent mechanisms and is strong enough to retain HaloTag containing an SV40 nuclear localization signal within the cytoplasm (Figure S3E). As expected, both CRS-Halo-RNase L and Halo-RNase L appeared restricted to the cytoplasm (Figure 3C-E). Furthermore, both constructs reduced EU pulse-labeled nascent RNA levels to a similar extent upon poly(I:C) transfection (Figure 3F-I), indicating that cytoplasmic RNase L is sufficient to reduce nascent RNA in the nucleus. In contrast, cells overexpressing CRS-Halo-NLS, a negative control, did not exhibit this pattern (Figure 3F, I). Taken together, these data suggest that activated RNase L does not induce nuclear RNA decay but rather reduces nascent RNA by affecting transcription.

### RNase L disrupts nucleoli and reduces serine-5-phosphorylated (S5P) Pol II levels in the nucleus

We first took an imaging-based approach to investigate whether RNase L activity affects the abundance or localization of transcriptional machinery. Pol I resides in the fibrillar component of nucleoli, which is surrounded by the nucleophosmin (NPM1)-containing granular component. Using antibodies against the large POLR1A subunit of Pol I and NPM1, we confirmed that POLR1A formed large aggregates surrounded by a ring-like pattern of NPM1 in mock-treated WT and RLKO cells (Figure 4A, Video S1-2). While the overall abundance of POLR1A and NPM1 did not significantly change upon Poly(I:C) treatment (Figure S4A-B), POLR1A aggregates and NPM1 rings became more dispersed in many of the WT cells, but less so in the RLKO cells (Figure 4A, yellow arrow denotes a WT cell exemplifying this phenotype, Video S3-4). We quantified this dispersion by calculating the coefficient of variation (defined as CV=σ/μ, where σ is the standard deviation and μ is the mean intensity) of POLR1A and NPM1 signals in the nucleus, with a lower CV indicating a more even (i.e., dispersed) distribution of signals. This confirmed a significantly reduced mean CV for both POLR1A and NPM1 in poly(I:C)-treated WT cells compared to RLKO cells (Figure 4B-C). Thus, consistent with the reduced Pol I-mediated rRNA transcription observed by 4sU-seq, RNase L activation leads to nucleolar disruption.

**Figure 4.**
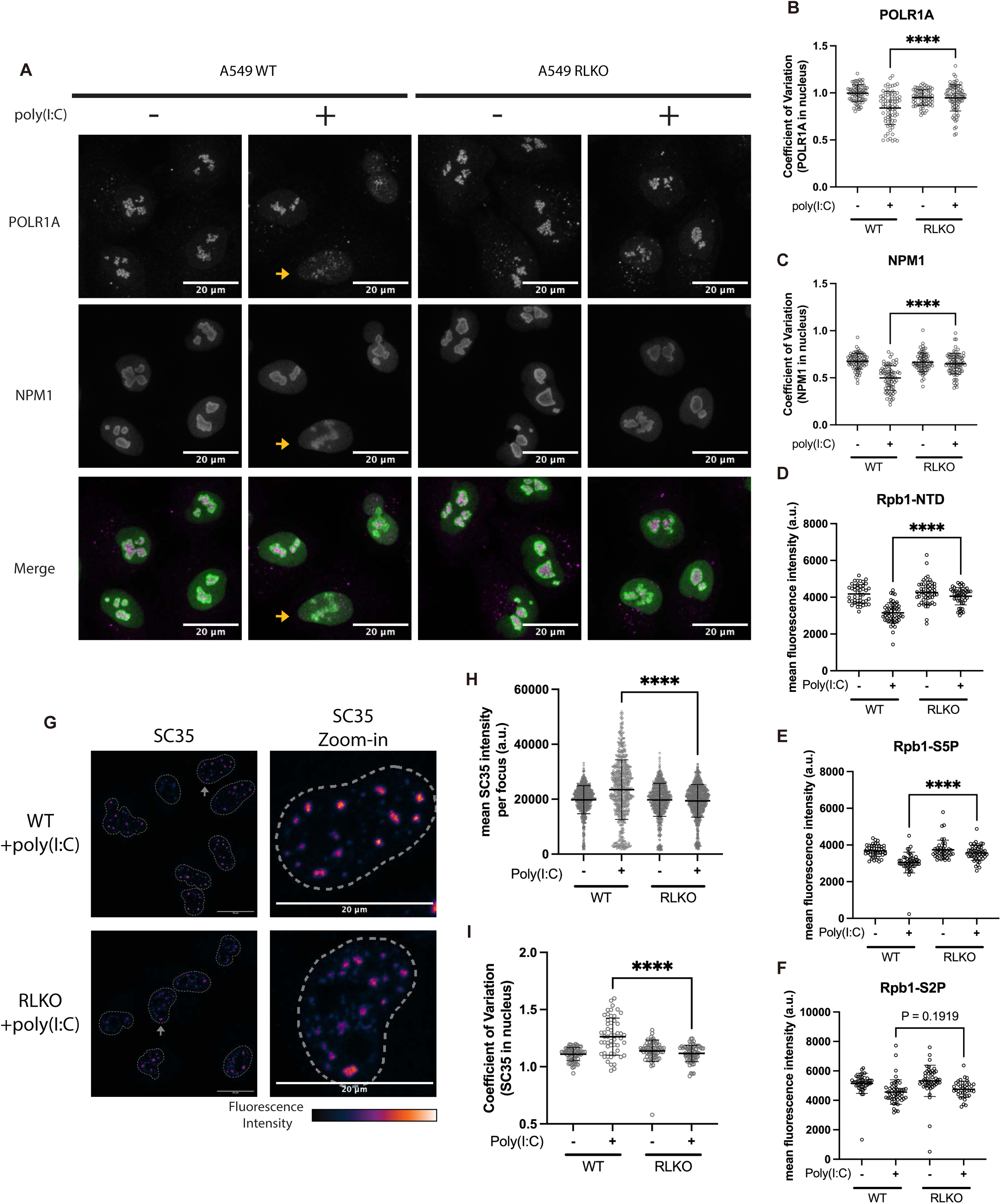
RNase L disperses nucleoli and reduces serine-5-phosphorylated Pol II levels in the nucleus. (A) Max-intensity-z-projection images showing POLR1A and NPM1 localization in poly(I:C)-transfected A549 WT or RLKO cells. Yellow arrows point to examples of nuclei showing the dispersal phenotype. (B-C) Quantification of the coefficient of variation (CV) of POLR1A (*B*) or NPM1 (*C*) in the nucleus, defined as CV=σ/μ, where σ is the standard deviation, and μ is the mean intensity. (D-F) Quantification of mean Pol II (NTD) (*D*), S5P (*E*), and S2P (*F*) staining intensity in poly(I:C)-transfected A549 WT or RLKO cells. (G) Max-intensity-z-projection images of SC35 localization in poly(I:C)-transfected A549 WT or RLKO cells. (H-I) Quantification of mean SC35 intensity for each SC35 focus (*H*) and the coefficient of variation of SC35 in the nucleus (*I*), using data in *(D)*. In all graphs, each point represents the value calculated for each nucleus. Error bars represent mean ± SD. ****, p<0.0001 (Welch’s t-test). The imaging data shown is representative of 3 independent biological replicates.

We next monitored the large RPB1 subunit of Pol II, which undergoes progressive phosphorylation within the heptad repeats in its C-terminal domain (CTD). We used antibodies recognizing either the RPB1 N-terminus (NTD, all phosphorylation states), the CTD serine 5 phosphorylated (S5P) version associated with transcription initiation, and the CTD serine 2 phosphorylated (S2P) version associated with elongation. Poly(I:C) treatment caused a modest but significant decrease in the abundance of total (NTD) and S5P Pol II in WT but not RLKO cells (Figure 4D-E, Figure S4C). We further validated the reduction of S5P Pol II by using an independent antibody (3E8) (Figure S4D). We did not observe consistent changes to total or phosphorylated Pol II localization, or to the abundance of S2P Pol II upon poly(I:C) treatment (Figure 4F).

Finally, we analyzed nuclear speckles, which are hubs for Pol II co-transcriptional RNA processing and undergo morphological changes in transcriptionally inhibited cells ^37^. Although nuclear speckle size was unchanged (Figure S4E), there was an increase in SC35 intensity and coefficient of variation in speckles in poly(I:C)-treated WT but not RLKO cells (Figure 4G-I, Figure S4F) as well as a more rounded speckle morphology (Figure S4G). Thus, RNase L activation is associated with a modest reduction in the abundance of unphosphorylated and S5P Pol II, as well as morphological changes to nuclear speckles and nucleoli.

### RNase L represses transcription via reduction of Pol II occupancy in the gene body

To directly measure how RNase L activation influences Pol II occupancy genome-wide, we performed CUT&RUN-sequencing using antibodies to the Pol II S5P CTD (4H8) and Pol II S2P CTD (E1Z3G) and applied yeast spike-in DNA normalization to account for potential global changes in occupancy. S5P and S2P Pol II occupancy were not globally changed at the transcription start site (TSS) in poly(I:C)-treated WT compared to RLKO cells or untreated cells (Figure 5A-B). Analyzing subnucleosomal fragments (<120bp) and nucleosomal fragments (>120bp) separately yielded similar results (Figure S5A-B), as did a differential binding analysis using DiffBind to measure Pol II occupancy on a gene-by-gene basis (Figure 5C-D).

**Figure 5.**
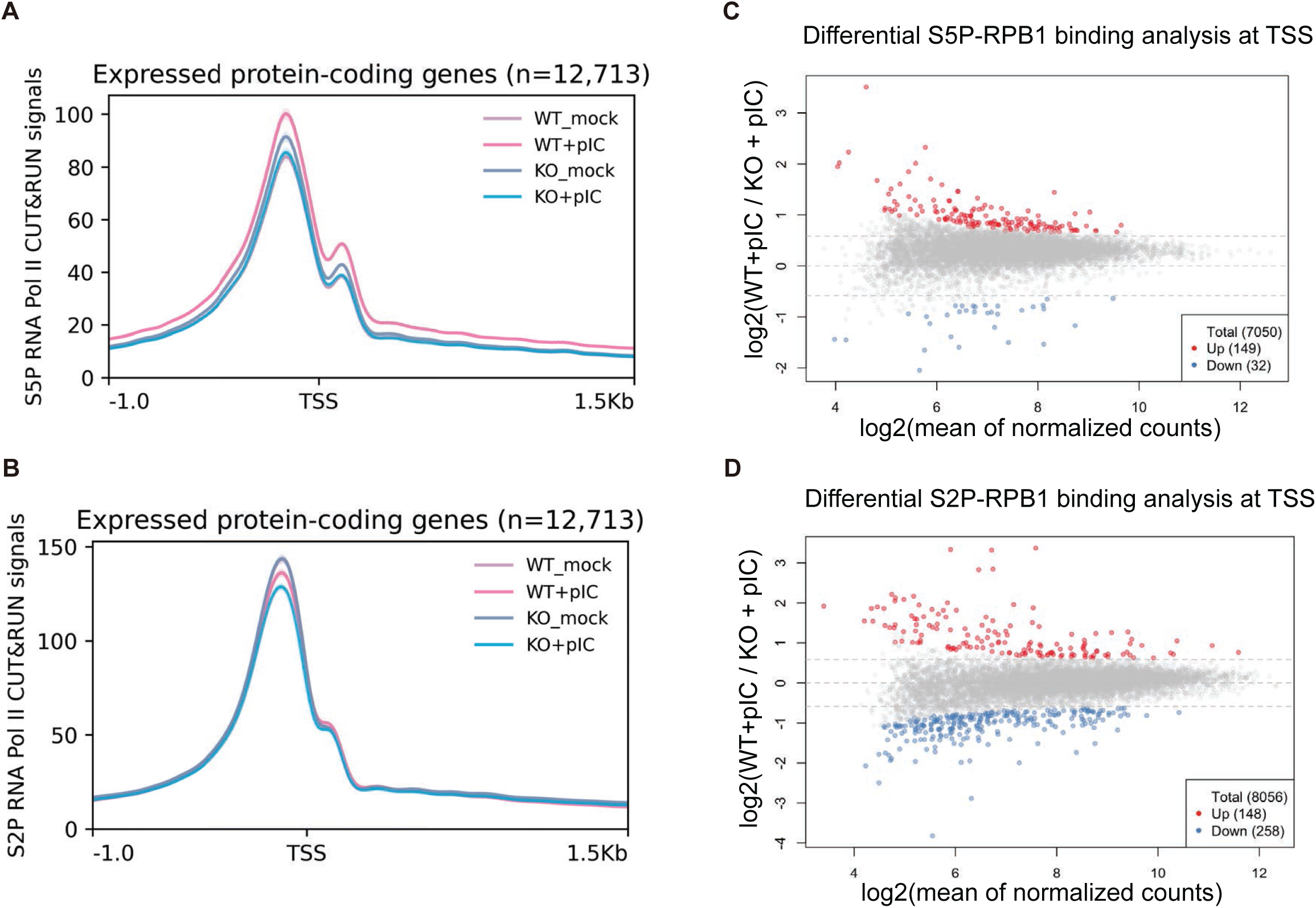
RNase L activation does not broadly inhibit Pol II occupancy at transcription start sites. (A-B) Metagene analysis of S5P (*A*) and S2P (*B*) Pol II occupancy at -1kb to +1kb around transcription start site (TSS) on expressed protein-coding genes in WT or RLKO cells, either mock-transfected or transfected with poly(I:C). Lines represent the average coverage of 3 independent biological replicates. (C-D) Differential binding analysis of S5P (*C*) and S2P (*D*) Pol II CUT&RUN-seq signal at the TSS (-500bp to +250bp) comparing WT cells transfected with poly(I:C) to RLKO cells transfected with poly(I:C). Significantly-up (red) and significantly-down (blue) are defined as adjusted *p*-value < 0.05 and log2(fold change) > 0.585 or < -0.585, respectively. Dashed lines denote log2(fold change) of -0.585, 0, and 0.585.

In contrast to the TSS occupancy data, when we zoomed in on the gene body and transcription end site (TES) regions, we observed a reduction in S2P Pol II occupancy (Figure 6A). To quantify this reduction, we first calculated an elongation index, defined as the density of reads in the gene body (+250bp from TSS to TES) divided by the density of reads at the TSS (-500bp to +250bp from TSS) (Figure 6B), with a lower elongation index suggestive of a defect of S2P Pol II traveling into the gene body. We observed an overall reduced elongation index in poly(I:C)-treated WT cells compared to poly(I:C)-treated RLKO cells or mock-treated cells (Figure 6C, S6A-B). This observation was further supported by a differential binding analysis across windows spanning different gene regions, which showed a progressive reduction of S2P Pol II binding midway through the gene body to the TES (Figure 6D-E). Approximately 30% of expressed genes show reduced S2P Pol II binding at the late-elongation window and the TES window (Figure 6F-G), and these also have a more pronounced reduction in 4sU RNA level (Figure S6C-D). Example traces are shown in Figure 6H, demonstrating reduced S2P occupancy within the gene body but no reduced S2P or S5P occupancy at the TSS of poly(I:C) treated WT relative to RLKO cells. While it is possible that the loss of S2P Pol II signal is due to reduced phosphorylation or enhanced dephosphorylation, such reduced S2P mark is also indicative of affected Pol II elongation, given the pivotal roles of S2P mark in regulating Pol II elongation ^38–40^. Thus, RNase L activation negatively impacts Pol II transcription primarily during the elongation phase.

**Figure 6.**
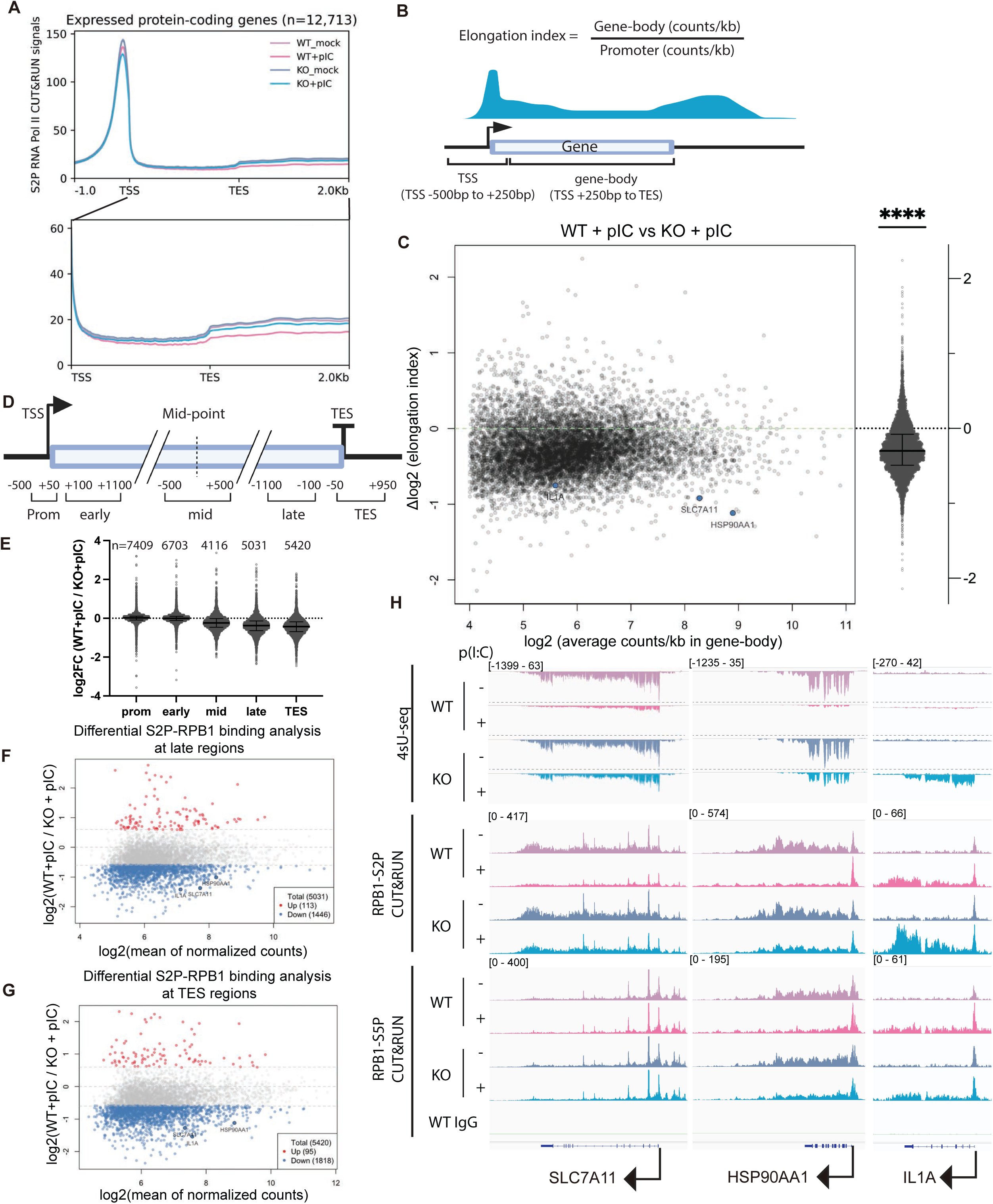
RNase L activation reduces S2P RNA Pol II occupancy in the gene body. (A) Metagene analysis of S2P Pol II occupancy on expressed protein-coding genes in WT or RLKO cells, either mock-transfected or transfected with poly(I:C). The lower panel represents a zoom-in view of coverage from the TSS to 2Kb after TES. Lines represent the average coverage of 3 independent biological replicates. (B) Diagram depicting the calculation of the elongation index and the range of the TSS window and the gene-body window. (C) Scatterplot showing the changes in the elongation index in relation to the read density in the gene body. The median and interquartile range of the changes in elongation index are shown to the right. ****, p < 0.0001 using one-sample t-test, comparing to 0. (D) Diagram depicting the range of promoter (Prom), early-elongation (early), mid-elongation (mid), late-elongation (late), and TES windows. (E) Summary of log2 fold changes comparing WT to RLKO cells, both transfected with poly(I:C), at windows defined in (*D*). Each dot represents the log2 fold change of S2P binding at the corresponding window within a gene, calculated by diffBind. Only genes longer than 3.2kb were included to minimize overlap. Number n represents quantifiable loci retained by diffBind after filtering. Error bars represent the median and interquartile range. (F-G) Zoom in view of the differential binding analysis in(E) of S2P Pol II CUT&RUN signal at the late elongation windows (-1100 bp to -100 bp from TES) (*F*) or TES windows (-50 bp to +950 bp from TES) (*G*). Significantly-up (red) and significantly-down (blue) are defined as adjusted *p*-value < 0.05 and log2(fold change) > 0.585 or < -0.585, respectively. Dashed lines denote log2(fold change) of -0.585, 0, and 0.585. (H) IGV tracks showing Pol II 4sU-seq, S2P CUT&RUN, and S5P CUT&RUN signal coverage on the housekeeping genes SLC7A11 and HSP90AA1 and the immune gene IL1A. All tracks represent the average coverage of 3 independent biological replicates. Coverage depth is shown in brackets in the upper track. For 4sU-seq tracks, a positive value indicating coverage in the positive strand direction, and a negative value indicating coverage in the negative strand direction.

### Importin β-mediated nuclear import links cytoplasmic RNA decay to transcription

Finally, we sought to determine the nature of the signal linking RNase L-induced cytoplasmic RNA decay with transcriptional repression in the nucleus. Accelerated cytoplasmic RNA decay is known to cause differential trafficking of RNA-binding proteins from the cytoplasm to the nucleus ^7,20^. To determine whether protein nuclear import plays a role in transcriptional repression by RNase L, we treated cells with Ibetazol, a specific inhibitor of importin β-mediated nuclear import ^41^, prior to measuring EU incorporation in our pulse labeling assay. Indeed, Ibetazol caused a significant increase in EU incorporation in poly(I:C)-treated WT cells but did not alter EU incorporation in RLKO cells (Figure 7A). Importantly, RNase L retained its mRNA degradation activity in Ibetazol-treated cells, as measured by RT-qPCR for RPLP0 and CHMP2A (Figure 7B). Thus, nuclear import of proteins, potentially those released during cytoplasmic RNA decay by RNase L, is required to induce subsequent transcriptional repression.

**Figure 7.**
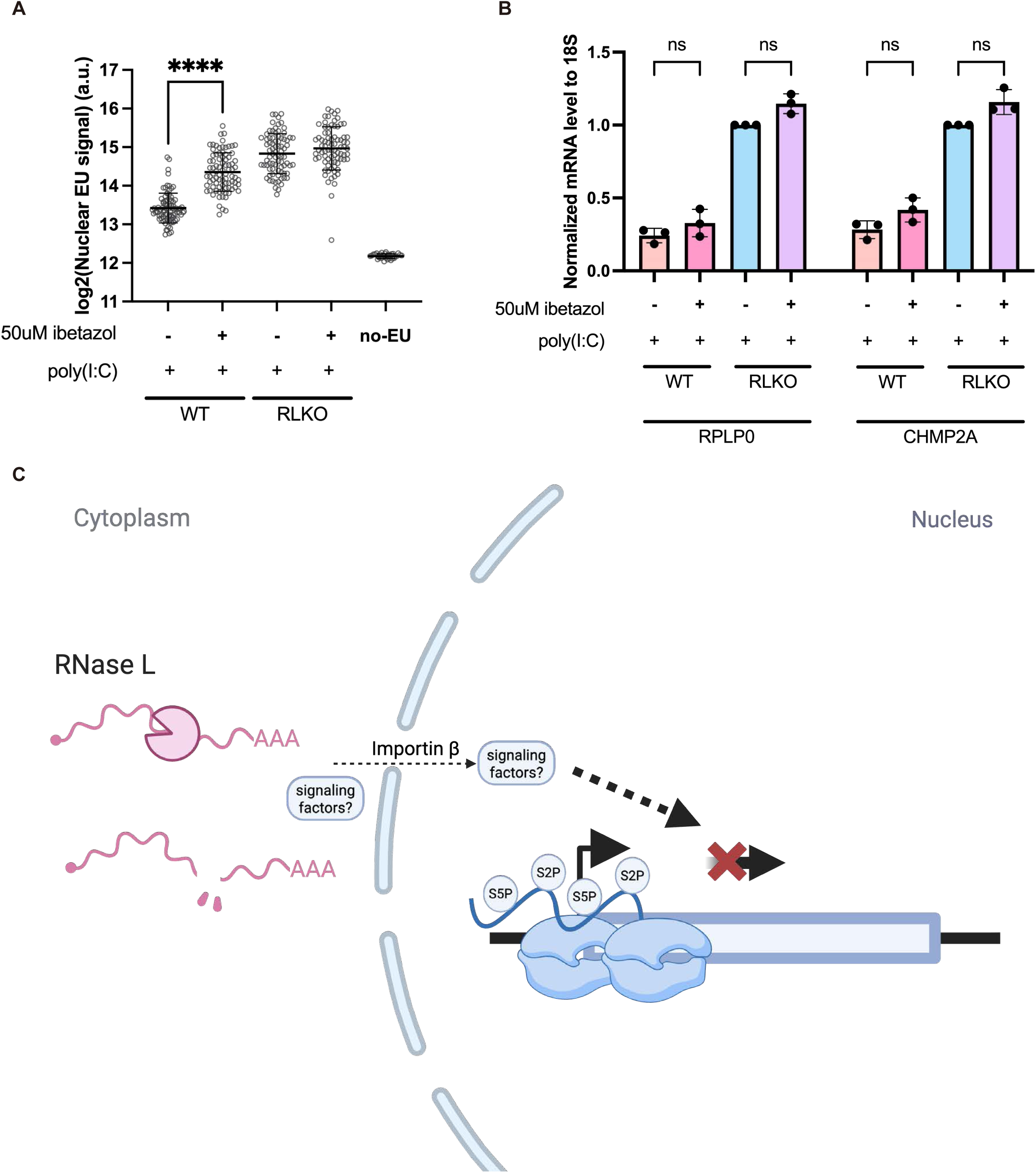
Inhibition of importin-β dampens transcription repression by RNase L. (A) Quantification of nuclear EU signal in WT or RLKO cells with indicated treatments. The data represent one representative replicate of four independent biological replicates. Each dot represents the mean nuclear EU fluorescence intensity of one individual nucleus. Bars represent mean ± SD. P-value was calculated using Welch’s *t* test. (B) RT-qPCR measuring the indicated housekeeping mRNA after poly(I:C) transfection in WT or RLKO cells, with or without ibetazol. Ns, not significant (p > 0.05 calculated with two-sided Welch’s *t* test). (C) Diagram depicting a model of RNase L-dependent transcription repression.

## Discussion

RNase L-induced cytoplasmic mRNA degradation extensively remodels the cellular gene expression landscape, influencing how cells respond to viral infection. Though originally thought to mainly inhibit translation, it is now appreciated that RNase L triggers cascading impacts on upstream stages of gene expression, including mRNA processing and export ^6,7^. Here, we demonstrate through EU pulse labeling, nascent mRNA sequencing, and Pol II occupancy experiments that cytoplasmic RNase L activity also profoundly impacts RNA synthesis by repressing transcription, including at immune genes. RNase L-induced transcriptional repression coincides with nucleolar disruption, as well as changes in nuclear speckle morphology, similar to those previously reported in transcriptionally inhibited cells^37^. Decay-to-transcription signaling involves protein shuttling between the two compartments, as inhibition of importin β breaks this connection and restores RNA synthesis (Figure 7C). The combination of these disruptions to the whole cellular gene expression pathway, spanning RNA synthesis through translation and decay, is likely what renders RNase L such a potent modulator of gene regulation.

We hypothesize that RNase L affects nascent mRNA primarily by impairing transcription elongation, as the most prominent Pol II phenotype was reduced S2P Pol II occupancy at the 3’ end of genes. However, it is possible that other stages of RNA biogenesis may also be impacted, especially given our imaging data showing a modest reduction in Pol II levels within the nucleoplasm. The fact that we observed reduced S5P Pol II levels by immunofluorescence but not by CUT&RUN suggests that a proportion of free or unstably bound S5P Pol II may undergo degradation. This could, for example, be mediated by the ARMC5 E3 ligase if the Pol II is defective ^42–44^. It is possible that, over time, a gradual reduction in total Pol II could lead to additional effects at earlier stages of transcription, or that reduced S5P Pol II selectively rather than globally impacts early-stage transcription in RNase L-activated cells.

Our observation that RNase L activation leads to changes to Pol I, NPM1, and SC35 distribution is consistent with previous reports that inhibition of Pol I and Pol II transcription can cause morphological changes to their respective transcription and processing sites ^45–47^. Notably, the dispersion of Pol I and NPM1 was similarly reported in cells treated with the cyclin-dependent kinase (CDK) inhibitor flavopiridol, and potentially due to inhibition of the phosphorylation of Pol I co-activator Treacle ^45^. Nuclear speckle-associated proteins play roles in co-transcriptional RNA processing. Similar to Pol II inhibition by chemical treatment, RNase L activation also altered nuclear speckle morphology, suggesting a link between transcription repression to other co-transcriptional defects, including intron retention ^7^ and reduced export ^6,46^ in RNase L-activated cells.

The fact that RNase L promotes transcriptional repression from the cytoplasm indicates that its mechanism of action is indirect and requires one or more signals to be conveyed between the compartments. We hypothesize this involves the relocalization of RNA-binding proteins (RBPs) from the cytoplasm to the nucleus, given that blocking protein nuclear import significantly increases RNA synthesis in RNase L-activated cells. More than 20 RBPs shuttle into the nucleus upon their release from mRNA undergoing degradation by herpesviral nucleases ^20^, and several similarly relocalize in RNase L-activated cells ^7^. Nuclear accumulation of these RBPs alters co-transcriptional RNA processing, including promoting mRNA hyperadenylation and nuclear poly(A) RNA accumulation ^6,7,15,48^. It is also associated with transcriptional repression by herpesviral nucleases ^19–21^. Thus, one possibility is that the accumulation of these RBPs in the nucleus disrupts co-transcriptional processing or the function of transcriptional regulators, leading to the observed elongation defects.

Other signals, in addition to protein redistribution, could also contribute to transcriptional repression by RNase L. For example, it could be propagated by stress signals conveyed by the global disruption of protein translation, e.g., as a consequence of ZAKα-dependent ribotoxic stress responses or other ribosome quality control pathways induced by RNase L ^8,11^. In support of this hypothesis, nascent protein ubiquitinylation in response to disrupted translation during heat shock contributes to transcriptional repression via a mechanism involving p38 signaling ^49^. Additionally, RNase L-induced translation arrest could result in a loss of proteins involved in global transcription regulation, particularly those with shorter half-lives. Consistent with this possibility, translation inhibition by cycloheximide in mouse embryonic stem cells can lead to depletion of euchromatin modifiers like Chd1 and reduced euchromatin marks, reduced Pol II occupancy, and less nascent RNA ^50^. However, our observation that nascent RNA synthesis was reduced as early as 2 hours post poly(I:C) transfection indicates that only highly labile proteins should be affected. Finally, it is possible that RNA fragments generated by RNase L cleavage serve as signaling molecules in the cytoplasm or upon transit into the nucleus ^51^.

A key outcome of transcriptional repression is the modulation of immune gene transcription. We find that although ISGs are robustly induced by poly(I:C) transfection independently of RNase L, RNase L activity tempers their induction. Many ISGs are downregulated due to reduced IFN signal amplification, which is relevant considering that RNase L activation is not uniform in cultures of cells exposed to dsRNA, and thus decreased gene expression by RNase L should influence innate immune signaling in neighboring cells ^4,6,52^. However, there are immune genes, such as IL1A, whose transcription is directly repressed by cytoplasmic RNA decay, indicating that RNase L-induced transcriptional repression can have both cell-autonomous and paracrine effects. Given the clinical implications of OAS and RNase L mutations in autoimmune diseases, including MIS-C ^1,9^, elucidating how the RNA decay-driven transcription repression shapes the immune landscape — in coordination with other branches of RNase L-driven pathways — should remain a focus of future studies.

### Limitations of the study

While our data support a defect in Pol II elongation in response to RNase L activation, we cannot rule out additional possible defects in other stages of transcription, including changes in CTD phosphorylation, initiation and pause release. We also do not know whether the transcriptional defects observed for Pol I and Pol III occur via a similar or distinct mechanism from that observed for Pol II. Finally, while we hypothesize that RBPs that are released from degraded RNA in the cytoplasm mediate the decay-to-transcription signaling during RNase L activation, an important future goal is to determine which specific proteins are required for transcriptional repression.

## Resource availability

### Lead contact

Further information and requests for resources and reagents should be directed to and will be fulfilled by the Lead Contact, Britt Glaunsinger.

### Materials availability

Plasmids generated in this study have been deposited to Addgene (see Key resources table for accession numbers).

## DATA AND SOFTWARE AVAILABILITY

4sU-seq, S5P Pol II CUT&RUN-seq, and S2P Pol II CUT&RUN-seq datasets were deposited in NCBI Gene Expression Omnibus under the accession numbers GSE298174, GSE298175, and GSE313712, respectively. No original code was reported. Further information required for reanalysis of data reported here will be available upon request.

## Acknowledgements

We thank all members of the Glaunsinger and Coscoy labs for helpful discussions. We thank James Burke and Roy Parker (CU Boulder) for providing the A549 RNase L KO and complementation cell lines and Susan Carpenter (UC Santa Cruz) for providing the iBMDM cell line. We thank Nharae Eugene Lee for generating the viral stocks used in this study. We thank Alejandro Rivera-Madera for generating the pLJM1-Halo-NLS plasmid used in this study. Imaging was performed at CRL Molecular Imaging Center (RRID:SCR_017852) and the Biological Imaging Facility at UC Berkeley, with special thanks to Holly Aaron and Denise Schichnes for training and technical support. Sequencing of the 4sU-seq samples were performed at Vincent J. Coates Genomics Sequencing Laboratory at UC Berkeley (QB3 Genomics, UC Berkeley, Berkeley, CA, RRID:SCR_022170). Sequencing of the CUT&RUN samples were performed at DNA Technologies and Expression Analysis Core at the UC Davis Genome Center, supported by NIH Shared Instrumentation Grant 1S10OD010786-01. B.G. is an investigator of the Howard Hughes Medical institute and this research was also supported by NIH R01CA136367. Diagrams were created with BioRender.

## Declaration of interests

The authors declare no competing interests.

## Figure legends

**Figure S1.**
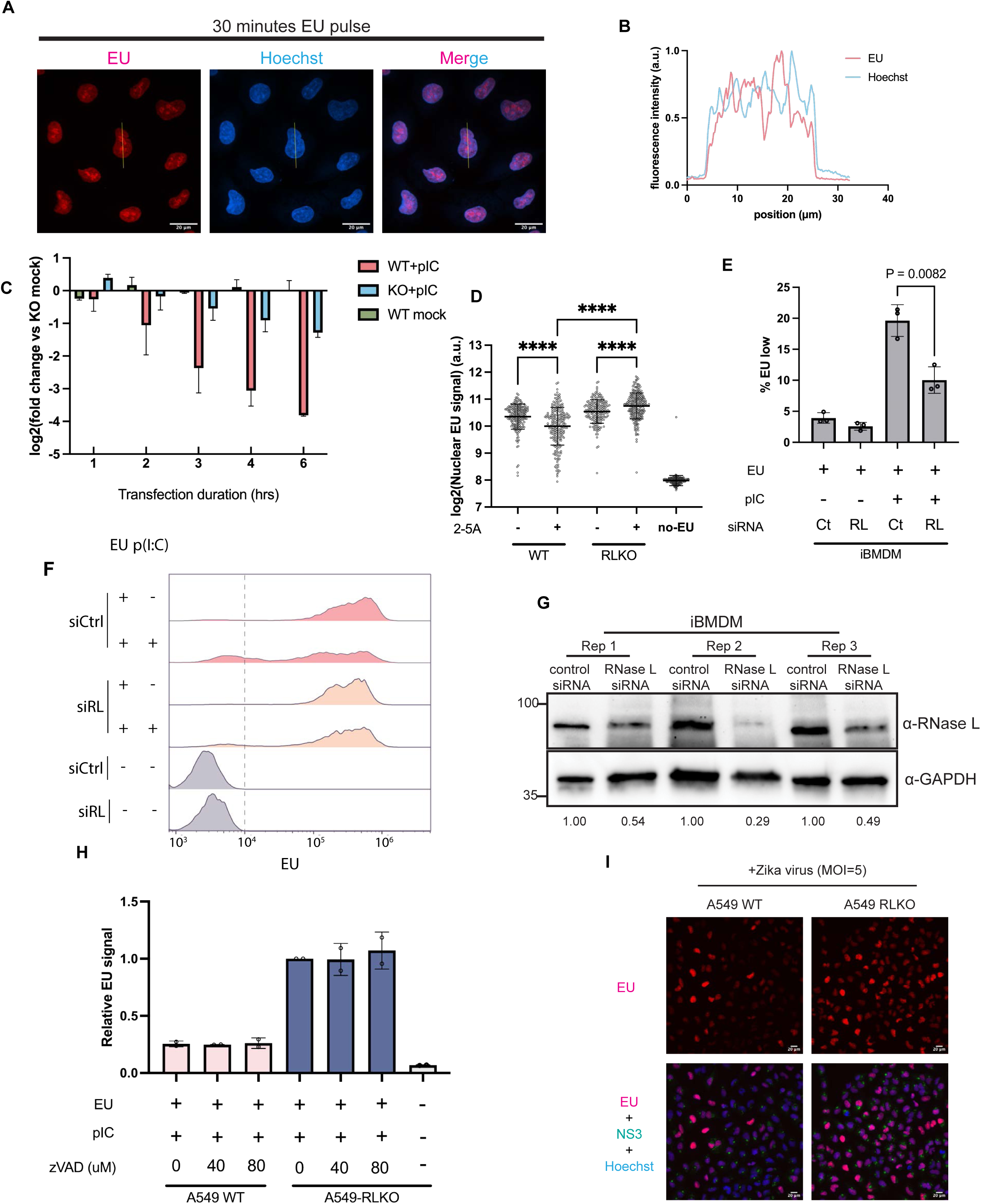
RNase L activation reduces nascent RNA progressively in A549 and iBMDM cells, independent of caspase activity. Related to Figure 1. (A) Representative max-intensity-projected images of mock-transfected A549 WT cells pulsed with 1mM EU for 30 minutes. (B) Normalized fluorescence intensity across line trace in (*A*). (C) A549 WT or RLKO cells were either mock-transfected with RNAiMAX or transfected with 0.76μg/ml poly(I:C) for durations indicated, followed by a 30-min pulse labeling with 1 mM EU. Error bars represent the mean ± SD of log2-fold change over mock-transfected RNase L KO cells at the indicated time point. Data is from 2 biological replicates. (D) Quantification of nuclear EU signal in WT or RLKO cells, either mock-transfected or transfected with synthetic 2-5A. The data show one representative replicate of two independent biological replicates. Each dot represents the mean nuclear EU fluorescence intensity of one individual nucleus. Bars represent mean ± SD. ****, p < 0.0001; P-value was calculated using two-sided Welch’s *t* test. (E) Quantification of the percentage of EU low cells. Error bars represent mean ± SD of 3 biological replicates. P-value was calculated using two-sided Welch’s *t* test. (F) A representative flow cytometry histogram of murine iBMDM cells with treatments indicated. EU signal was gated according to no-EU controls, shown with a dashed line. (G) Western blot validation of RNase L siRNA knockdown efficiency. Numbers below lanes indicate relative levels of bands normalized to the control KD of the corresponding replicate. (H) A549 WT or RNase L KO cells were transfected with 0.76μg/ml poly(I:C) in the presence of z-VAD at the indicated concentration for 4 hours, followed by 30-min pulse labeling with 1mM EU. Each dot represents the median fluorescence intensity of the sample with indicated treatment from one biological replicate, normalized to the median fluorescence intensity of DMSO-treated KO cells. Bars represent mean ± SD. (I) Low magnification (20X) field of view of Zika virus-infected cells, from the same sample as Figure 1D-E.

**Figure S2.**
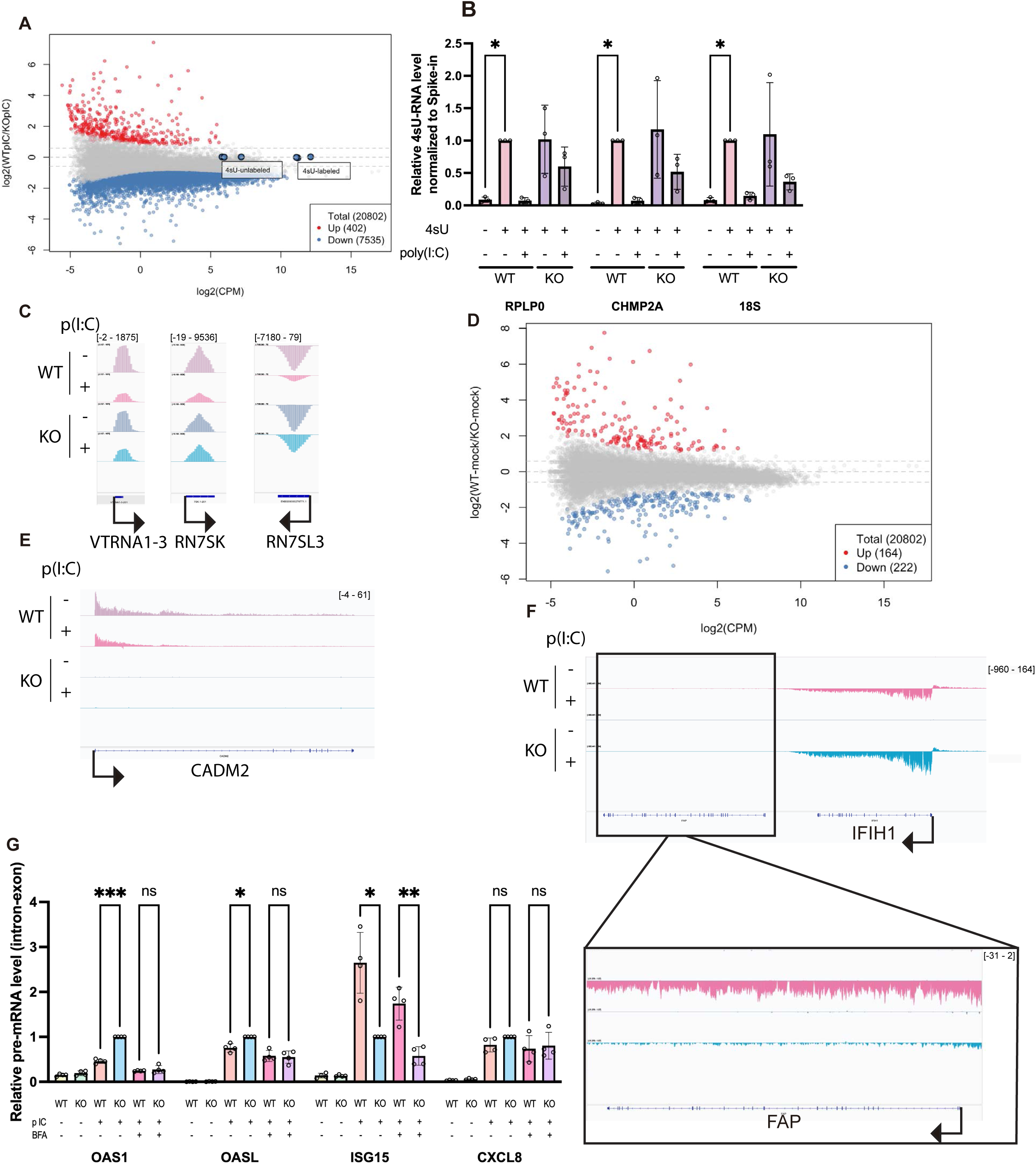
4sU-seq data accurately captures changes on the nascent RNA level. Related to Figure 2. (A) Differential expression analysis of 4sU-seq comparing WT cells to RLKO cells transfected with poly(I:C) cells (same as Figure 2D), with spike-ins highlighted. See Figure 2D for details. (B) RT-qPCR of 4sU samples compared to no-4sU control. RNA samples were the same as the ones used in 4sU-seq. *, p < 0.05 (ratio paired *t* test). (C) IGV tracks showing 4sU-seq signal coverage on RN7SK, VTRNA1-3 and RN7SL3, examples of downregulated Pol III transcribed genes. Tracks represent the average coverage of 3 independent biological replicates. In this and subsequent tracks, a positive value indicates coverage in the positive strand direction, while a negative value indicates coverage in the negative strand direction, and coverage depth is shown in brackets in the upper track. (D) Differential expression analysis of 4sU-seq comparing mock-transfected WT cells RLKO cells. (E) IGV tracks showing 4sU-seq signal coverage on CADM2, an example of cell line-specific expression variation. Tracks represent the average coverage of 3 independent biological replicates. (F) IGV tracks showing 4sU-seq signal coverage on FAP downstream of IFIH1, an example of read-through transcription. Tracks represent the average coverage of 3 independent biological replicates. (G) Pre-mRNA level of additional immune genes in WT or RLKO cells transfected with poly(I:C), with or without treatment of brefeldin A (BFA). Pre-mRNA level is determined by RT-qPCR using intron-exon spanning primer pairs, normalized to 18S. *, p< 0.05; **, p<0.01; ***, p< 0.001; ns, not significant; calculated with two-sided Welch’s *t* test.

**Figure S3.**
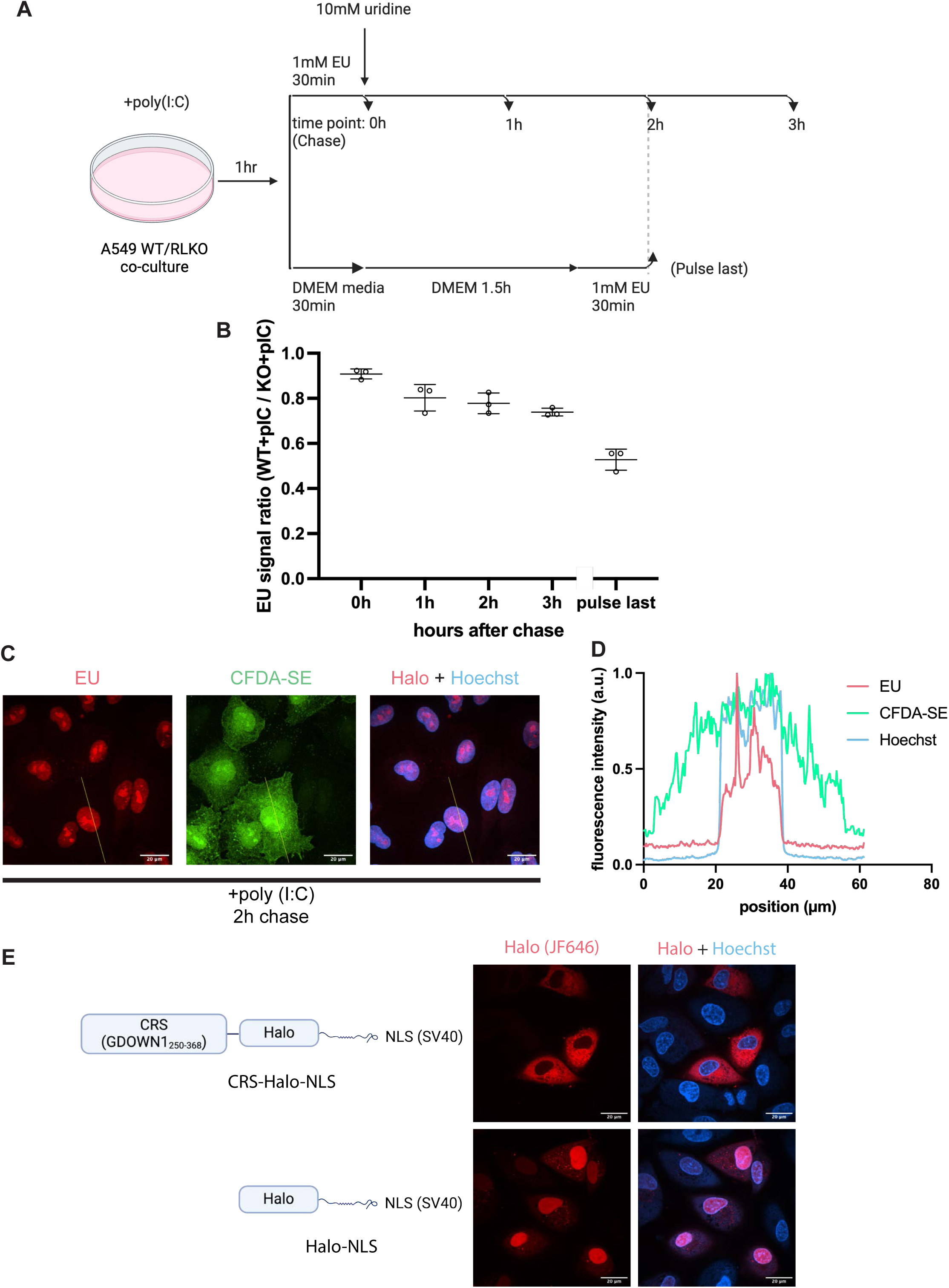
EU pulse chase experiments accurately measure nuclear RNA turnover kinetics and functional confirmation of the CRS tag. Related to Figure 3. (A) Diagram showing the experimental setup of the “EU pulse last” control in parallel to the EU pulse chase experiments shown in Figure 3A. (B) Quantification of EU signal in the “EU pulse at last” control. Data from Figure 3B, replotted as the ratio of EU signals in A549 WT to RLKO cells, are included for comparison. Each dot represents one biological replicate. Bars represent mean ± SD. (C) Representative max-intensity-projected images of cells transfected with poly(I:C) for 1 hour, pulsed with 1mM EU for 30 minutes then chased for 2 hours. WT cells were labeled with CFDA-SE dye and co-cultured with RLKO cells. (D) Normalized fluorescence intensity across line trace in *(C)*. CFDA-SE signal is included as a uniform distributed control for comparison. (E) Representative images of the localization of CRS-Halo-NLS and Halo-NLS. A single z plane is shown per construct. Diagram showing the constructs are included.

**Figure S4.**
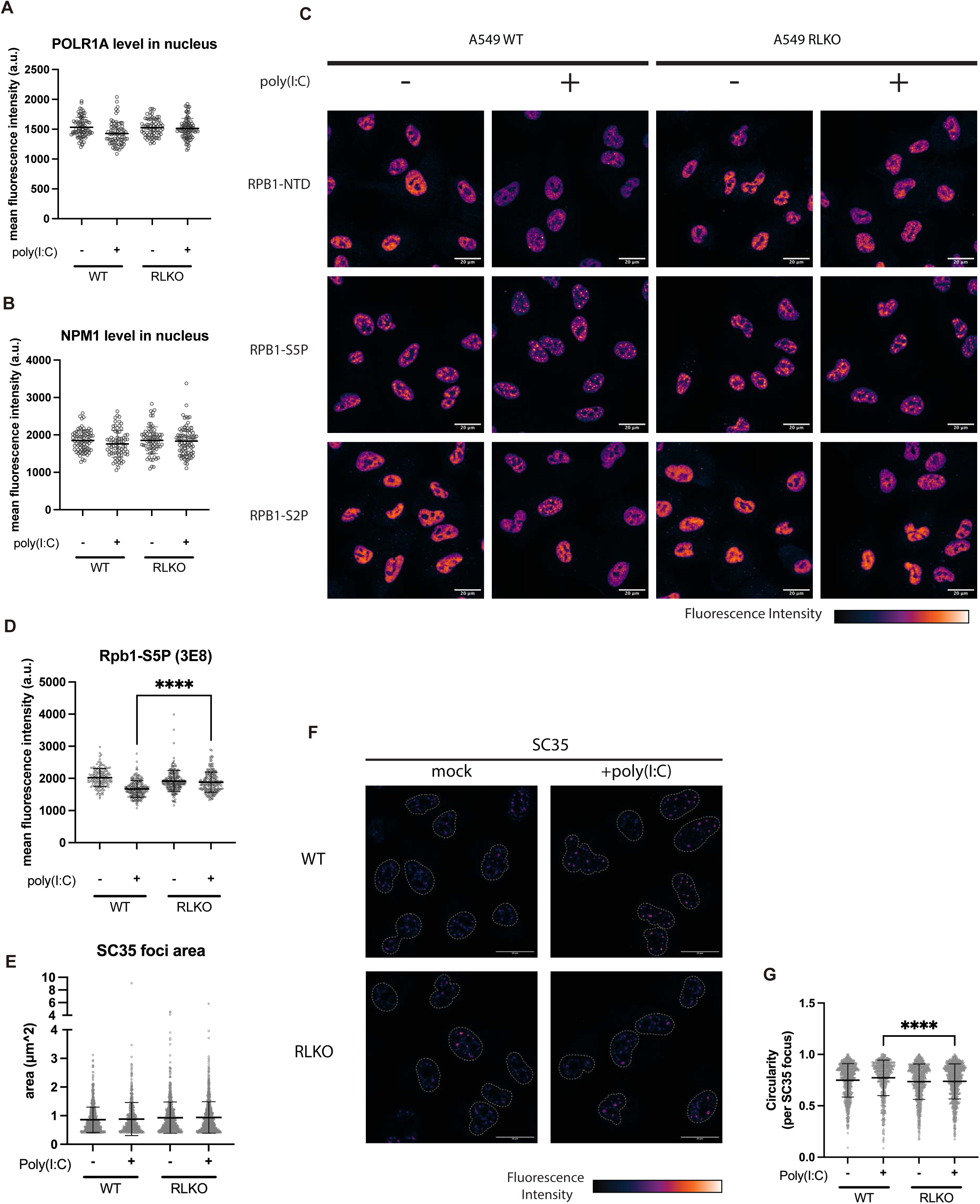
RNase L activation reduces total and S5P Rpb1 levels in the nucleus and does not aYect nuclear speckle size. Related to Figure 4. (A-B) Quantification of mean POLR1A (A) and NPM1 (B) staining intensity in poly(I:C)-transfected A549 WT or RLKO cells. (C) Representative max-intensity-z-projection images of Pol II (NTD), S5P, and S2P staining in A549 WT or RLKO cells transfected as indicated. Data are the same as Figure 4D-F. (D) Quantification of mean S5P-RPB1 staining intensity using 3E8 antibody in poly(I:C)-transfected A549 WT or RLKO cells. Each point represents the mean intensity calculated for each nucleus. Error bars represent mean ± SD. ****, p<0.0001 (Welch’s t-test). Data represents 1 replicate of 2 independent biological replicates. (E) Quantification of SC35 foci area for each SC35 focus using data in Figure 4D. Each point represents the value measured for each SC35 focus. Error bars represent mean ± SD. Data are from 1 representative replicate of 3 total replicates. (F) Max-intensity-z-projection images of SC35 localization in mock or poly(I:C)-transfected A549 WT or RLKO cells. Data from Figure 4G is included for comparison. (G) Quantification of mean SC35 circularity for each SC35 focus using data in Figure 4G and Figure S4F. Each point represents the value calculated for each SC35 focus. Error bars represent mean ± SD. ****, p<0.0001 (two-sided Welch’s t-test).

**Figure S5.**
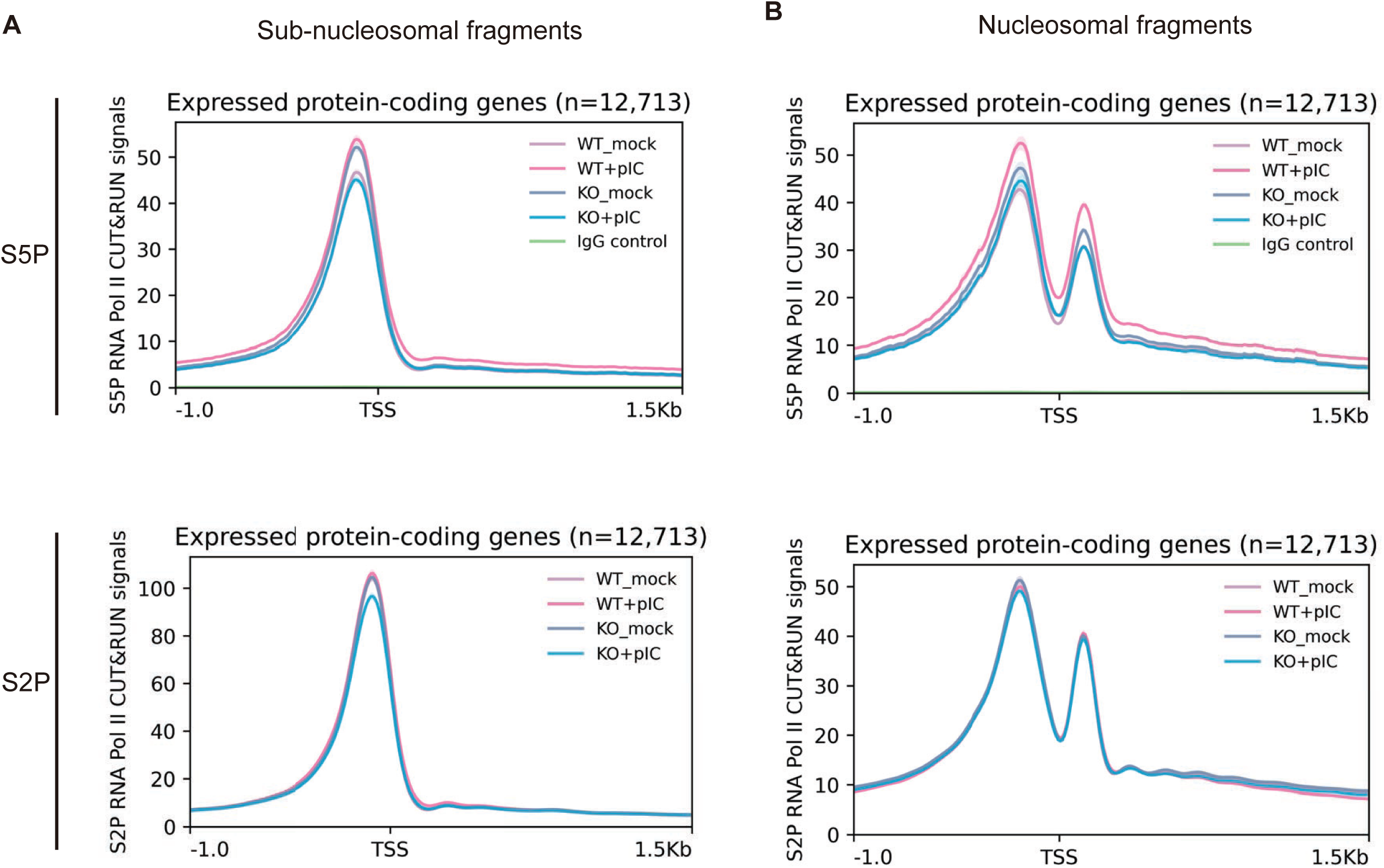
RNase L activation does not reduce CUT&RUN subnucleosomal and nucleosomal signals at TSS. Related to Figure 5. (A) Metagene analysis of S5P and S2P Pol II CUT&RUN subnucleosomal fragment (<120bp) coverage at -1kb to +1.5kb around transcription start site (TSS) of expressed protein-coding genes. (B) Metagene analysis of S5P and S2P Pol II CUT&RUN nucleosomal fragment (>120bp) coverage at -1kb to +1.5kb around the transcription start site (TSS) of expressed protein-coding genes. The enrichment of subnucleosomal fragments slightly upstream of TSS and the bimodal peaks around the TSS of the nucleosomal fragments likely reflect the local nucleosome organization around Pol II occupancy sites.

**Figure S6.**
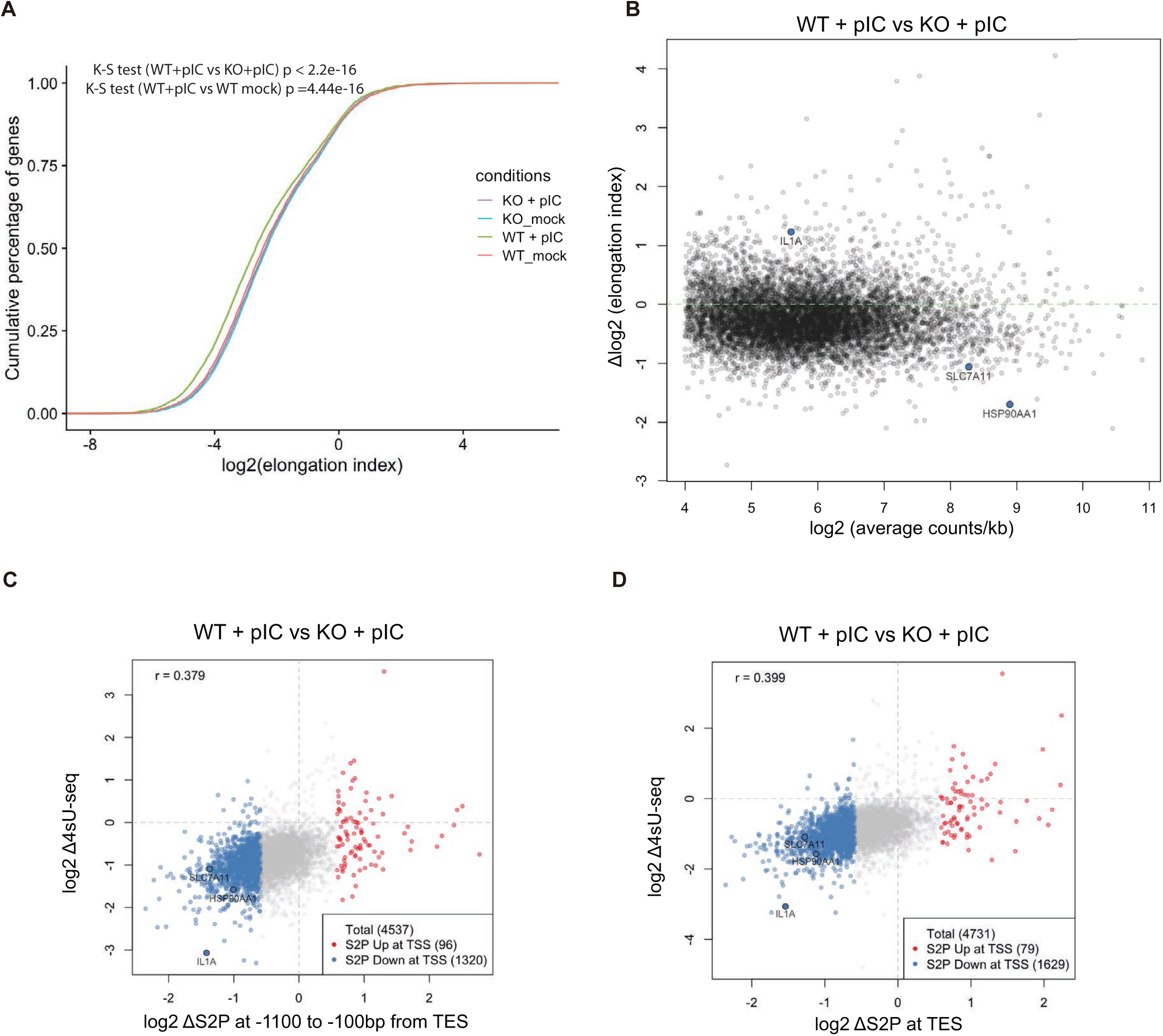
RNase L activation reduces S2P Pol II CUT&RUN signal in the gene body, which correlates with the reduction in 4sU-seq signal. Related to Figure 6. (A) Empirical cumulative distribution function (ECDF) plot of log2(elongation factor) for WT or RLKO cells, either mock-transfected or transfected with poly(I:C). P values were calculated with two-sided Kolmogorov–Smirnov tests. (B) Scatterplot showing the changes in elongation index between poly(I:C)-transfected WT cells and mock-transfected WT cells in relation to the read density in the gene body. (C-D) Scatterplot showing the relation between log2 fold change of 4sU-seq and S2P Pol II occupancy at late (*C*) or TES windows (*D*). Up- or down-regulated S2P is defined in the same way as Figure 6F-G.

## STAR METHODS

### Key resources table

| REAGENT or RESOURCE | SOURCE | IDENTIFIER |
| --- | --- | --- |
| **Antibodies** |  |  |
| mouse anti-NS3 antibody (Clone E1D8) | Balsitis et al., 2009<sup>53</sup> | N/A |
| POLR1A (D6S6S) Rabbit mAb | Cell Signaling Technology | Cat# 24799;<br>RRID:AB_2798884 |
| Nucleophosmin antibody [FC82291] | Abcam | Cat# ab10530;<br>RRID:AB_297271 |
| Rpb1 NTD (D8L4Y) | Cell Signaling Technology | Cat# 14958;<br>RRID:AB_2687876 |
| Rpb1 S5P CTD (ab5131) (for immunofluorescence) | Abcam | Cat# ab5131,<br>RRID:AB_449369 |
| Rpb1 S5P CTD (3E8) (for immunofluorescence) | Active Motif | Cat# 61085,<br>RRID:AB_2687451 |
| Rpb1 S5P CTD (4H8) (for CUT&RUN) | Cell Signaling Technology | Cat# 2629,<br>RRID:AB_2167468 |
| Rpb1 S2P CTD (E1Z3G) (for CUT&RUN) | Cell Signaling Technology | Cat# 13499,<br>RRID:AB_2798238 |
| Rpb1 S2P CTD (ab5095) (for immunofluorescence) | Abcam | Cat# ab5095,<br>RRID:AB_304749 |
| Mouse Anti-SC35 Monoclonal Antibody | Abcam | Cat# ab11826,<br>RRID:AB_298608 |
| Recombinant Anti-RNase L antibody [EPR15894] | Abcam | Cat# ab191392,<br>RRID:AB_2916269 |
| GAPDH antibody [6C5] | Abcam | Cat# ab8245,<br>RRID:AB_2107448 |
| goat anti-rabbit IgG AlexaFluor Plus 488 | Thermo Fisher Scientific | Cat# A32723,<br>RRID:AB_2633275 |
| goat anti-mouse IgG AlexaFluor 546 | Thermo Fisher Scientific | Cat# A-11035,<br>RRID:AB_2534093 |
| donkey anti-rat IgG AlexaFluor Plus 488 | Thermo Fisher Scientific | Cat# A48269,<br>RRID:AB_2893137 |
| goat anti-rabbit IgG AlexaFluor 647 | Thermo Fisher Scientific | Cat# A27040,<br>RRID:AB_2536101 |
| Goat Anti-Rabbit IgG-HRP | SouthernBiotech | Cat# 4030-05,<br>RRID:AB_2687483 |
| Mouse normal IgG | Santa Cruz Biotechnology | Cat# sc-2025,<br>RRID:AB_737182 |
| **Bacterial and virus strains** |  |  |
| Zika virus (Nica 2–16) | Tabata et al., 2016<sup>54</sup> | GenBank accession<br>KX421194.1 |
| **Chemicals, peptides, and recombinant proteins** |  |  |
| 4-thiouridine (4SU) | Sigma Aldrich | T4509-100MG |
| HPDP-Biotin | Thermo Fisher Scientific | PI21341 |
| pAG-MNase | Cell Signaling | 40366S |
| proteinase K | Cell Signaling | 10012S |
| RNase A | Thermo Fisher Scientific | EN0531 |
| 4-Thio-UTP | Jena Bioscience | NU-1156S |
| T7 RNA polymerase | NEB | M0251L (25,000U) |
| CFDA-SE | Cayman Chemical | 14456 |
| 5-Ethynyl-uridine | Lumiprobe | 2439-500mg |
| Sulfo-Cyanine5-Azide | Lumiprobe | A3330 |
| Alexa Fluor 488 Azide | Thermo Fisher Scientific | A10266 |
| Ibetazol | Axon MedChem | Axon 4300 |
| In-Fusion® snap assembly (master mix) | Clontech (TaKaRa) | 638943 |
| TURBO DNase | Thermo Fisher Scientific | AM2239 (5,000U) |
| AMV Reverse Transcriptase | Promega | M5101 (300U) |
| cOmplete EDTA-free protease inhibitor cocktail | Roche | 5056489001 |
| Halt Protease and Phosphatase Inhibitor Cocktail (100X) | Thermo Fisher Scientific | 78440 |
| Poly(I:C) (HMW) | InVivoGen | tlrl-pic |
| Lipofectamine RNAiMAX | Thermo Fisher Scientific | 13778150 |
| Brefeldin A Solution (1000X) | Thermo Fisher Scientific | 00-4506-51 |
| Janelia Fluor 646 HaloTag Ligand | Promega | GA1121 |
| Critical commercial assays |  |  |
| KAPA Stranded RNA-Seq Kit with RiboErase (HMR) | Roche | KK8484<br>07962304001 |
| KAPA Single-Indexed Adapter Kit | Roche | KK8700<br>08005699001 |
| ERCC RNA Spike-In Mix | Thermo Fisher Scientific | 4456740 |
| NEBNext Ultra II DNA Library Prep Kit | NEB | cat # E7645L |
| NEBNext Multiplex Oligos for Illumina (Unique Dual Index UMI Adaptors DNA Set 1) | NEB | cat # E7395S |
| Universal Mycoplasma Detection Kit | ATCC | 50189644FP |
| iTaq universal SYBR green supermix | Bio-Rad | 1725121 |
| Q5 Hot Start High-Fidelity 2X Master Mix | NEB | M0494S |
| Deposited data |  |  |
| Raw and processed 4sU-seq data | This study | GEO: GSE298174 |
| Raw and processed CUT&RUN-seq data | This study | GEO: GSE298175 |
| Total RNA-seq for mock- or poly(I:C)-transfected A549 cells | Burke et al., 2019<sup>2</sup> | GEO: GSE124144 |
| Experimental models: Cell lines |  |  |
| A549 WT cells | Burke et al., 2019<sup>2</sup> | N/A |
| A549 RLKO cells | Burke et al., 2019<sup>2</sup> | N/A |
| A549 RLKO cells reconstituted with 3xflag-RNase L (A549 RLKO-RL) | Burke et al., 2019<sup>2</sup> | N/A |
| A549 RLKO cells reconstituted with catalytic dead 3xflag-RNase L (R667A) (A549 RLKO-CM) | Burke et al., 2019<sup>2</sup> | N/A |
| Murine immortal bone-marrow-derived macrophages (iBMDM) | Robinson et al., 2021<sup>55</sup> | N/A |
| Oligonucleotides |  |  |
| 2-5A (pA<sub>(2-5A)</sub>)<sub>4</sub> | IDT | N/A |
| ON-TARGETplus Non-targeting Control Pool | Dharmacon | D-001810-10-05 |
| SMARTPool: ON-TARGETplus Mouse RNASEL siRNA, 10nM | Dharmacon | L-043480-00-0010 |
| Table S1 | This paper | N/A |
| Recombinant DNA |  |  |
| pLJM1-Halo-RNaseL | This study | Addgene: 250856 |
| pLJM1-CRS-Halo-RNaseL | This study | Addgene: 250857 |
| pLJM1-CRS-Halo-NLS | This study | Addgene: 250859 |
| pLJM1-Halo-NLS | This study | Addgene: 250858 |
| Software and algorithms |  |  |
| HTStream |  | RRID:SCR_018354 |
| STAR | Dobin et al., 2013<sup>56</sup> | RRID:SCR_004463 |
| FeatureCounts | Liao et al., 2014<sup>57</sup> | RRID:SCR_012919 |
| Limma-voom | Ritchie et al., 2015<sup>58</sup> | RRID:SCR_010943 |
| topGO | DOI: 10.18129/B9.bioc.topGO | RRID:SCR_014798 |
| Deeptools | Ramirez et al., 2016<sup>59</sup> | RRID:SCR_016366 |
| Trimmomatic | Bolger et al., 2014<sup>60</sup> | RRID:SCR_011848 |
| Bowtie 2 | Langmead and Salzberg., 2012<sup>61</sup> | RRID:SCR_016368 |
| SAMTOOLS | Li et al., 2009<sup>62</sup> | RRID:SCR_002105 |
| UMI-tools | Smith et al., 2017<sup>63</sup> | RRID:SCR_017048 |
| DiffBind | Ross-Innes et al., 2012<sup>64</sup> | RRID:SCR_012918 |
| FIJI | Schindelin et al., 2012<sup>65</sup> | RRID:SCR_002285 |
| ImageJ | Schneider et al., 2012<sup>66</sup> | RRID:SCR_003070 |
| Prism 10 v10.3.0 | GraphPad | N/A |
| R | www.r-project.org | N/A |
| RStudio | www.rstudio.com | N/A |
| Other |  |  |
| Dynabeads MyOne Streptavidin C1 | Thermo Fisher Scientific | 65002 |

## CONTACT FOR REAGENT AND RESOURCE SHARING

### Experimental model and subject details

A549 WT, RLKO, RLKO-RL and RLKO-CM cell lines ^2^ were generously provided by James Burke (University of Florida Scripps) and Roy Parker (University of Colorado, Boulder). Immortalized murine bone marrow macrophages ^55^ were generously provided by Susan Carpenter (UC Santa Cruz). All cell lines were maintained in DMEM supplemented with 10% FBS and tested for mycoplasma routinely using Universal Mycoplasma Detection Kit (ATCC, cat # 50189644FP).

## METHOD DETAILS

### Plasmids

All plasmids generated in this study were sequence verified by whole plasmid sequencing and have been deposited on Addgene, with accession numbers indicated in the Key Resources Table. All PCR reactions were performed using Q5 2X master mix (NEB, cat # M0494S). Sequences encoding the CRS tag, HALO tag, and RNase L with appropriate overhangs were PCR amplified. pLJM1-Halo-RNaseL was constructed by directly cloning the HALO and RNase L PCR fragments into linearized pLJM1-puro ^67^ using InFusion (Takara Bio, Cat # 639650). pLJM1-CRS-Halo-RNaseL was constructed by first assembling CRS, HALO and RNase L PCR fragments into a single fragment by overlap extension PCR, followed by cloning into linearized pLJM1-puro using InFusion. pLJM1-CRS-Halo-NLS was generated from pLJM1-CRS-Halo-RNaseL using a site-directed mutagenesis protocol described previously ^68^ to replace RNase L fragment with an SV40 NLS (PKKKRKV). pLJM1-Halo-NLS was generated from pLJM1-Halo-RNaseL using the same mutagenesis method as above.

### EU pulse chase for flow cytometry

Cells were were mock-transfected, transfected with 0.76μg/ml poly(I:C) HMW (InVivoGen, cat # tlrl-pic), or 10μM 2-5A (pA(_2-5_A)_4_, custom synthesized by IDT) using Lipofectamine RNAiMAX (Thermo Fisher, cat # 13778-150) as indicated, then the media was replaced with 1mM EU (Click Chemistry Tools cat #CCT-1261-500 or Lumiprobe cat # 2439-500mg) in DMEM with 10% FBS. The cells were incubated at 37°C for 30 minutes. For flow cytometry samples, cells were washed once with 1ml PBS, harvested, and fixed with 1% PFA. The cells were then permeabilized by resuspending in 100ul 0.05% w/v saponin in PBS. To visualize EU incorporation, 500ul click reaction mix (4mM CuSO_4_, 100mM sodium ascorbate, 2 μM sulfo-cyanine5-azide (Lumiprobe cat# A3330) in TBS, 50mM Tris-HCl, pH 8.0, 150mM NaCl) was added to the cell suspension and the samples were incubated at room temperature for 30 minutes, protected from light. The samples were then washed twice with 1ml 0.05% w/v saponin in PBS, once with 1ml PBS, then resuspended in PBS before flow cytometry analysis. The samples were then analyzed on an BD Accuri C6 Plus flow cytometer.

For EU pulse chase experiments, WT cells were first color-labeled with 1 μM CFDA-SE (Cayman Chemical, cat # 14456) in PBS for 10 min at room temperature, quenched with equal volume of DMEM with 10% FBS, then washed with DMEM with 10% FBS 3 times. The labeled WT cells were resuspended in DMEM with 10% FBS, pooled with unlabeled RLKO cells at a 1:1 ratio and were seeded in 6-well plates the day before experiment. Cells were transfected with 0.76μg/ml poly(I:C) HMW (InVivoGen, cat # tlrl-pic) using Lipofectamine RNAiMAX (Thermo Fisher, cat # 13778-150) for 1 hour and pulsed with 1mM EU in DMEM with 10% FBS for 30 minutes at 37°C. The EU-containing media was removed, and cells were washed with PBS once and immediately chased with 10mM uridine in DMEM with 10% FBS. The samples were harvested at the indicated time-points. An additional well without transfection and EU pulse labeling was harvested at the same time as the 0-hour chase samples, serving as a no-EU control. The “Pulse last” sample, which served as a control for transfection efficiency, was transfected with 0.76μg/ml poly(I:C) for 1 hour, incubated DMEM with 10% FBS without EU for 30 minutes at 37°C, washed with PBS once and replaced the supernatant with DMEM with 10% FBS and incubated for 1.5 hours. Then the cells were pulsed with 1mM EU in DMEM with 10% FBS for 30 minutes at 37°C and harvested at the same time as the 2-hour chase samples.

To characterize EU incorporation in cells overexpressing tagged RNase L, A549 RLKO cells were first nucleofected with the indicated construct using Neon transfection system (Invitrogen) according to the manufacturer’s recommended settings for A549 cells (1230V, 30ms, 2 pulses), using 5μg DNA per 750,000 cells in 100μL E2 buffer. The nucleofected cells were cultured in 6-well plates overnight. The cells were then incubated with 20nM Halo ligand Janelia Fluor 646 (Promega, cat # GA1121) diluted in 1mL DMEM + 10%FBS per well 30 minutes prior to poly(I:C) transfection, then transfected with 0.76μg/ml poly(I:C) for 4 hours without removing the Halo dye and pulsed with 1mM EU for 30 minutes same as above. The samples were processed as above, with the exception that 5uM Alexa Fluor™ 488 Azide (Thermo Fisher, cat #A10266) was used.

To characterize EU incorporation in murine iBMDM cells depleted of RNase L, cells were maintained in DMEM with 10% FBS and grown to 90% confluence. Cells were removed and washed once with Dulbecco’s phosphate-buffered saline (DPBS) (Gibco). Nucleofections were done using the Neon Transfection System (Thermo Fisher). 2 × 10^6^ cells were resuspended in 100 μL of buffer R, to which control non-targeting or RNase L targeting pools of siRNA (ON-TARGETplus SMARTpool siRNA – Horizon Discovery) were added to a final concentration of 200nM. This was loaded into a Neon 100-μL pipette tip and Neon tube with 3 mL of buffer E2 with electroporation parameters set to 1,680 V, 20 ms, 1 pulse. Following electroporation, cells were plated in 10 mL of DMEM with 10% FBS in 10 cm TC-treated plates and were incubated at 37°C for 24 hours. Cells were then transfected with 0.76μg/ml poly(I:C) pulsed with 1mM EU for 30 minutes same as above. The samples were processed as above.

### Immunofluorescence

For Zika infection in Figure 1D-E and Figure S1G, A549 WT or RLKO cells were seeded in glass-bottomed 24 well-plates at a density of 100,000 cells per well the day before the experiment, and after media change, were infected with Zika virus (Nica 2–16, GenBank accession KX421194.1) at an MOI = 5 for 24 hours. Cells were treated with 1 mM EU for 30 minutes before being washed once with 1 mL of PBS, then fixed with 4% PFA for 15 minutes. The cells were then permeabilized with 0.5% Triton X-100 in PBS for 15 minutes and washed 3 times with PBS. Samples were then incubated with 500 μL of click reaction mix at room temperature for 30 minutes, followed by two washes with PBS and storage in PBS. Samples were then blocked with 1% BSA in PBS with 0.02% Tween-20 for 1 hour and incubated with mouse anti-NS3 antibody (Clone E1D8) diluted 1:100 in the blocking solution (Balsitis et al., 2009). The samples were then washed three times with PBS + 0.02% Tween-20, incubated with goat anti-mouse IgG AlexaFluor Plus 488 (Invitrogen, cat #A32723) diluted 1:1000 in blocking solution, and washed three times with PBS + 0.02% Tween-20. The nuclei were stained with 8 μM Hoechst 33342 in PBS for 30 minutes, followed by 2 PBS washes.

For RNase L localization studies, A549 RLKO cells were nucleofected as in the “EU pulse chase for flow cytometry” section and 340,000 cells were seeded per well in a glass-bottomed 24-well plate and incubated overnight. Cells were first labeled with 20nM Halo ligand Janelia Fluor 646 for 30 minutes and transfected with poly(I:C) at a concentration of 0.76μg/ml for 4 hours without removing the Halo ligand. The cells were then pulse labeled with EU and fixed, permeabilized, labeled with click reaction and Hoechst 33342 as above.

For characterizing changes in POLR1A, NPM1, RPB1 and nuclear speckle morphology, A549 WT or RLKO cells were seeded in glass-bottomed 24 well-plates at a density of 100,000 cells per well the day before experiment and transfected with either lipofectamine RNAiMAX alone or with poly(I:C) at a concentration of 0.76μg/ml for 4 (optionally pulsed with 1mM EU for 30min) to 4.5 hours. Samples fixed and permeabilized and optionally labeled with click reaction as above, then were blocked with 0.3%-0.5% BSA in PBS with 0.3-0.5% Triton X-100 for 0.5-1 hour. Then samples were stained with primary antibodies against POLR1A (clone D6S6S, Cell Signaling, #24799, 1:500), NPM1 (Abcam, ab10530-100, 1:500), Rpb1 NTD (clone D8L4Y, Cell Signaling Technology, 1:500), Rpb1 S5P (ab5131, Abcam, 1:500; or 3E8, Active Motif, RRID:AB_2687451), Rpb1 S2P (ab5095, Abcam, 1:500) and SC35 (ab11826, Abcam, 1:500) diluted in blocking solution overnight at 4°C, goat anti-rabbit IgG AlexaFluor Plus 488 (Invitrogen, cat #A32723) alone or goat anti-mouse IgG AlexaFluor 546 (Invitrogen, cat # A11035), donkey anti-rat IgG AlexaFluor Plus 488 (Invitrogen, cat # A48269), and goat anti-rabbit IgG AlexaFluor 647 (Invitrogen, cat # A27040) diluted 1:1000 in blocking solution as secondaries, and 8-10μM Hoechst 33342.

For Ibetazol treatment, A549 WT or RLKO cells were seeded in glass-bottomed 24 plates at a density of 100,000 cells per well the day before the experiment and transfected with poly(I:C) at a concentration of 0.76μg/ml for 3 hours in the presence of DMSO or 50 μM Ibetazol. Cells were then pulse labeled with EU and fixed, permeabilized, labeled with click reaction.

All samples were imaged on a Nikon Eclipse Ti spinning disk confocal microscope or a Zeiss LSM 880 laser scanning microscope.

### Image processing

All confocal image processing was performed using FIJI ^65^, either manually or with custom macros. Line profile measurements were performed using the plot profile function, and the data were exported to Prism 10 for normalization and visualization. For nuclear signal quantification in Figure 1D-E and Figure 4D-F, the nuclear regions of interest (ROIs) were generated from the Hoechst channel in the image z-stack by Gaussian blur, thresholding with the Otsu method, and 3D segmentation using the 3D Manager. To fill holes within ROIs and exclude objects on edges, the 3D segmentation map was converted to binary, followed by “3D fill holes” and “3D exclude edges”. The processed binary map was 3D segmented again using the 3D Manager to generate the final ROIs. The ROIs were then used to quantify mean intensity in selected channels with the “quantify” function in 3D Manager.

For quantification in Figure 4B-C, H-I, 7A, and S4D, the image z-stack was first max-intensity or sum-intensity projected. Then the nuclear ROIs were generated from the Hoechst channel by Gaussian blur, thresholding with the Otsu method, hole filling by the “Fill holes” function, and size selected with the “Analyze particle” function to only include particles larger than 50 μm ^2^. Then the ROIs were used to quantify mean intensity and standard deviation in selected channels with the “Measure” function. For nuclear speckle analysis, a binary mask for the nuclei was first generated, then the channels of interest were multiplied to the mask to isolate nuclei. A duplicated SC35 channel was then used to generate nuclear speckle ROIs by Gaussian blur, local thresholding with the Bernsen method, size selected with the “Analyze particle” function to only include particles larger than 0.4 μm^2^. Then the ROIs were used to quantify mean intensity, area, and circularity in original SC35 channel with the “Measure” function.

### 4sU-spike-in RNA production using in-vitro transcription

4sU-labeled or unlabeled spike-in RNA was produced according to a protocol described previously with modifications ^69^. Briefly, ERCC RNA Spike-In Mix (Thermo Fisher, 4456740) was reverse transcribed using AMV-RT (Promega, cat # M5108) and random 9-mer primer (IDT). Amplicons containing the T7 promoter were amplified using primers in Table S1 and Q5 2X master mix. In vitro transcription was performed using T7 RNA polymerase (NEB, M0251L) and 5 mM each of ATP, GTP, and CTP. For labeled spike-in, 4.5mM UTP and 0.5mM 4-thio-UTP (Jena Bioscience, NU-1156S) were used; for unlabeled spike-in, 5mM UTP was used. Template DNA was then digested with TURBO DNase. RNA was then purified by phenol:chloroform extraction and cleaned up with Zymo RNA clean and concentrator -5 to remove unincorporated 4-thio-UTP. An equal mass of 3 labeled spike-in RNAs and 3 unlabeled spike-in RNAs was pooled and used for 4sU-seq.

### 4sU-Seq

Six million A549 WT or RLKO cells were seeded in 15cm tissue culture dishes and cultured overnight before transfection. Then, cells were then transfected with poly(I:C) for 4 hours, followed by pulse labeling with 500μM 4sU dissolved in DMEM media supplemented with 10% FBS for 10 minutes. The plates were washed once with PBS, then lysed with 2ml TRIzol. RNA was extracted from the TRIzol lysate, followed by isopropanol precipitation. 100μg or 300μg (WT with poly(I:C)) total RNA was combined with an equal amount of pooled spike-in RNA and was biotinylated with HPDP-biotin (Thermo Scientific cat # PI21341, 50μg per 25μg of RNA) for 2 hours, followed by two chloroform extractions and isopropanol precipitation. The RNA samples were denatured at 65°C for 5 minutes, then incubated with Dynabeads MyOne Streptavidin C1 magnetic beads (Thermo Fisher, cat # 65002) that had been previously blocked with glycogen for 1 hour. The beads were washed twice with wash buffer (100 mM Tris-HCl, 10 mM EDTA, 1 M sodium chloride, and 0.1% Tween 20) at 65°C then twice at room temperature. To increase the stringency of the washes, the beads were further washed with TE buffer (10mM Tris-HCl pH 7.4, 1mM EDTA) twice at 55°C ^70^. 4sU-RNA was then eluted from the beads twice with 100 μL of 100 mM DTT, and the combined eluent was precipitated with isopropanol.

The sequencing library was constructed using KAPA Stranded RNA-Seq Kit with RiboErase (HMR) (cat # KK8484, 07962304001) and KAPA Single-Indexed Adapter Kit (Roche, cat # KK8700, 08005699001) following manufacturer’s instructions, with the following modifications: Briefly, 60ng to 100ng RNA was used as input without rRNA depletion and fragmented at 85 °C for 6 minutes; the PCR cycle numbers were determined empirically by qPCR. The library was submitted to QB3-Berkeley Genomics Sequencing Core for QC and sequencing on a NovaSeq 6000 S1 chip.

### Analysis of published RNA-seq datasets

Published total RNA-seq datasets for mock- or poly(I:C)-transfected A549 cells (GEO: GSE124144) ^2^ were used to as a proxy to assign the dominant transcript for each gene. The raw fastq files were first pre-processed using HTStream (https://github.com/s4hts/HTStream) then pseudo-aligned and quantified using Kallisto ^71^. For each gene, the highest expressed transcript (measured by transcript-per-million, TPM, first criteria) with the longest transcript length (second criteria) was selected as the dominant transcript. Only genes on the main nuclear chromosomes (i.e., chr1-22, chrX and chrY) and the corresponding dominant transcript with average TPM > 0 are included.

### 4sU-Seq data processing

The raw reads were pre-processed using HTStream v1.3.2 (https://github.com/s4hts/HTStream), which included the separation of rRNA and non-rRNA reads into separate files via HTStream SeqScreener. The rRNA reads were aligned to the rDNA repeat (GenBank Accession U13369.1), concatenated with ERCC sequences, and the non-rRNA reads were aligned to the human genome (hg38), concatenated with ERCC sequences, using STAR v2.6.1b ^56^. When aligning rRNA reads, options *--alignIntronMax 1* and *--alignSJDBoverhangMin 999* were set to suppress gapped alignment. Aligned reads were counted using Rsubread (v2.16.1) FeatureCounts ^57^, and differential gene expression analysis was performed with limma-Voom v3.58.1, normalized to 4sU-labeled spike-in. ^58^. A model of ∼0+group+replicate is used to account for batch effects. Segments of pre-rRNA (i.e., 5’ ETS, 18S, ITS1, 5.8S, ITS2, 28S, 3’ ETS) were included in the differential expression analysis. Scaled, stranded BigWig files were generated with Deeptools bamCoverage function, using scaling factors generated in limma-Voom (proportional to the reciprocal of norm.factors × lib.sizes). BigWig files of biological replicates were averaged using Deeptools bigwigAverage function. For both operations, bin sizes were set to 20bp. The metagene coverage plots were generated using Deeptools ^59^ using the coordinates of dominant transcripts defined in “Analysis of published RNA-seq datasets”. GO term analysis was performed with the topGO package (DOI: 10.18129/B9.bioc.topGO), using the Weight01 algorithm and Fisher’s exact test.

### CUT&RUN-sequencing

The CUT&RUN procedure was performed according to previous reports ^72,73^, with modifications. Briefly, 250,000 A549 WT or RNase L KO cells were seeded per well in a 12-well plate and cultured overnight before transfection. Then cells were then transfected with RNAiMAX alone or with 0.76 μg/ml poly(I:C) for 4 hours The cells were permeabilize on plate with 500μL Perm buffer (20 mM HEPES-KOH pH 7.5, 150 mM NaCl, 0.5 mM Spermidine (Sigma, S2626), 0.1% Triton X-100 (Sigma, X100), and proteinase inhibitor (Roche, 04693132001)) for 15 minutes at room temperature. The cells were washed again with Perm buffer then incubated with either anti-Rpb1 S5P CTD (4H8, Cell Signaling, cat #2629) at 1: 50 dilution, equal mass of mouse normal IgG (Santa Cruz, cat # sc-2025), or anti-Rpb1 S2P CTD (clone E1Z3G, Cell Signaling, cat # 13499) at 1: 50 dilution, in 200μL Perm buffer for 1 hour at room temperature. The cells were then washed twice with Perm buffer and incubated with pAG-MNase (Cell Signaling, cat #40366) at a 1:33 dilution in 200μL Perm buffer for 1 hour at room temperature. The cells were then washed twice with Perm buffer and chilled on ice. Then, 225 μL ice-cold Perm buffer containing 5 mM CaCl_2_ was dispensed per well, and the plate was incubated in a refrigerator at 4°C for 30 min for targeted digestion. The reaction was halted by adding 75 μL 4× STOP solution (680 mM NaCl, 40 mM EDTA, 8 mM EGTA, 100 μg/ml RNase A (Invitrogen, cat # 12091021) and 0.1% Triton X-100, spiked with yeast genomic DNA (Cell Signaling, cat #40366) at 10pg per sample for 4H8 and IgG, 30pg per sample for E1Z3G). The plate was then incubated at 37°C for 30min to facilitate fragment release. The supernatant was collected, and then 3μL 20% SDS solution and 3.75μL 20mg/ml proteinase K solution were added. The mixture was then incubated at 50°C for 1 hour. The DNA was extracted with phenol:chloroform.

The sequencing library was constructed with NEBNext® Ultra™ II DNA Library Prep Kit (NEB, cat # E7645L) following a previous report ^74^ with modifications. Briefly, equal volume of purified DNA fragments was used. End preparation was performed at 20°C for 30 min and then at 50°C for 60 min to prevent short fragment melting. Fragments were then ligated to adaptors from NEBNext® Multiplex Oligos for Illumina® (Unique Dual Index UMI Adaptors DNA Set 1, NEB, cat # E7395S) and cleaned up twice with 1.1X Ampure XP beads (Beckman Coutler, cat # A63880). The resulting ligated products were then PCR amplified for 10 cycles for 4H8 and E1Z3G, and 14 cycles for IgG samples, respectively. The cycle numbers were determined empirically by qPCR. The amplified libraries were double-sided size-selected with 0.6X and 1.1X Ampure XP beads and further cleaned up by 1 round with 1.1X beads for 4H8, E1Z3G, or 3 rounds for IgG samples, to remove adaptor dimers. The libraries were sequenced on AVITI platform (Element Biosciences) by UC Davis sequencing core.

### CUT&RUN data processing

Reads were trimmed with Trimmomatic v0.39 ^60^ and aligned to a concatenated human (hg38) and yeast (SacCer3) genome with bowtie 2 v2.5.2 ^61^. The BAM files were then deduplicated with UMI_tools v1.1.5 ^63^ and optionally splitted into subnucleosomal (<120bp) and nucleosomal (>120bp) fractions using Deeptools (v3.5.5) alignmentSieve. The spike-in normalization factors were calculated by counting reads mapped to yeast genome using Samtools idxstats^62^, followed by conversion to scaling factors that are proportional to the reciprocal to the spike-in read counts. BigWig files were generated with Deeptools bamCoverage function, using scaling factors generated above. BigWig files of biological replicates were averaged using Deeptools bigwigAverage function. For both operations, bin sizes were set to 20bp. The metagene coverage plots were generated using Deeptools. Differential binding analysis was performed with DiffBind v3.12.0 ^64^ using the built-in spike-in normalization function in *dba.normalize*. A model of ∼0+group+replicate is used to account for batch effects. For analysis in Figure 6, different windows were analyzed separately to avoid merging of regions by diffBind. For elongation index calculation, FeatureCounts was used to count reads mapped to either TSS (-500bp to +250bp) or gene body (+250bp to TES). One pseudocount was then added to all regions. Reads were then normalized using the spike-in scaling factors generated above. Only genes with >16 counts/kb at both the TSS and the gene body were kept for subsequent calculation. Elongation index was averaged by taking the geometric mean of 3 replicates, followed by logarithmic conversion.

### Quantitative reverse transcription PCR (RT-qPCR)

500,000 A549 WT or RLKO cells per well were seeded in 6-well plates the day before either mock-transfecting or transfecting with 0.76μg/ml poly(I:C), in the absence or in the presence of 1X brefeldin A (eBiosciences, cat # 00-4506-51) when indicated, for 3 hours. RNA samples were prepared by TRIzol extraction. They were then treated with Turbo DNase (Thermo Fisher, cat # AM2238) for 30 minutes at 37°C and re-purified by phenol:chloroform extraction. 500ng to 1μg RNA was used for reverse transcription using AMV-RT (Promega, cat # M5108) and random 9-mer primer (IDT). cDNA was then quantified by qPCR using iTaq Universal SYBR Mastermix (Bio-Rad, cat # 1725125) and the indicated primer pairs. Transcript levels were normalized to 18S.

For Figure S2B, an equal volume of 4sU RNA eluent was directly reverse-transcribed using AMV-RT (Promega) and a random 9-mer primer (IDT), and quantified as above with the indicated primer pairs. Transcript levels were normalized to spike-in.

For Figure 7B, 1μg RNA was treated with Turbo DNase for 15 minutes at 37°C and inactivated at 70°C in the presence of 15mM EDTA. The treated RNA was diluted with DEPC water to 50μL and 4μL was used for reverse transcription using AMV-RT and random 9-mer primer. cDNA was then quantified by qPCR with iTaq Universal SYBR Mastermix.

### Western blotting

To prepare whole-cell lysates for evaluating RNase L knock-down, murine iBMDMs transfected with RNase L targeting siRNAs (Horizon, cat # L-043480-00-0010) were washed with cold DPBS (Gibco) followed by lysis with radioimmunoprecipitation assay (RIPA) lysis buffer (50 mM Tris HCl, 150 mM NaCl, 1.0% [vol/vol] NP-40, 0.5% [wt/vol] sodium deoxycholate, 1.0 mM EDTA, and 1% [wt/vol] SDS, Halt™ Protease and Phosphatase Inhibitor Cocktail (Thermo Fisher, cat # PI78440). Cell lysates were vortexed briefly, rotated at 4°C for 15 min, and then clarified by centrifugation at 21,000 × *g* in a tabletop centrifuge at 4°C for 10 min to remove debris. 30 μg of whole-cell lysate were resolved on 4% to 15% mini-PROTEAN TGX gels (Bio-Rad). Transfers to polyvinylidene difluoride (PVDF) membranes (Bio-Rad) were done with the Trans-Blot Turbo transfer system (Bio-Rad). Blots were incubated in 5% milk in TBS with 0.1% Tween 20 (TBS-T) to block, followed by incubation with primary antibodies against anti-RNase L (Abcam, ab191392, 1:10,000) and anti-GAPDH antibody (abcam ab8245, 1;5,000). Washes were carried out with TBS-T. Blots were then incubated with HRP-conjugated secondary antibodies (Southern Biotechnology, 1:5,000). Washed blots were incubated with Clarity Western ECL substrate (Bio-Rad) for 5 min and visualized with a ChemiDoc imager (Bio-Rad). Band intensity was analyzed with ImageJ ^66^.

## QUANTIFICATION AND STATISTICAL ANALYSIS

All statistical analyses were performed with GraphPad Prism 10 and RStudio. For limma-Voom analysis for 4sU-seq, and DiffBind analysis for CUT&RUN-seq, adjusted P value or False discovery rate (FDR) is corrected with the Benjamini-Hochberg (BH) method. All statistical method details and the p value notation are included in figure legends.

**Video S1-S4. Related to Figure 4**. Z-stack movies showing the localization of POLR1A (left panel) and NPM1 (center panel) or the merged view (right panel) in mock-treated WT (*video S1*) or RLKO (*video S2*) A549 cells or p(I:C)-treated WT (*video S3*) or RLKO (*video S4*) A549 cells.

**Supplemental Table 1.** Primer sequences used in this study.

## Notes

### Competing Interest Statement

The authors have declared no competing interest.

### Summary of Updates

New data on Pol I localization was included. Results on Pol II localization was modified. Related discussion was modified. Current version included new dataset of S2P Pol II CUT&RUN. Spike-in normalization was used for all CUT&RUN data analysis instead. Related discussion was modified.

